# Neurospectrum: A Geometric and Topological Deep Learning Framework for Uncovering Spatiotemporal Signatures in Neural Activity

**DOI:** 10.1101/2023.03.22.533807

**Authors:** Dhananjay Bhaskar, Yanlei Zhang, Jessica Moore, Feng Gao, Bastian Rieck, Guy Wolf, Firas Khasawneh, Elizabeth Munch, J. Adam Noah, Helen Pushkarskaya, Christopher Pittenger, Valentina Greco, Smita Krishnaswamy

**Author notes:** Co-first authors (Dhananjay Bhaskar, Yanlei Zhang, Jessica Moore).

## Abstract

Neural signals are high-dimensional, noisy, and dynamic, making it challenging to extract interpretable features linked to behavior or disease. We introduce *Neurospectrum*, a framework that encodes neural activity as latent trajectories shaped by spatial and temporal structure. At each timepoint, signals are represented on a graph capturing spatial relationships, with a learnable attention mechanism highlighting important regions. These are embedded using graph wavelets and passed through a manifold-regularized autoencoder that preserves temporal geometry. The resulting latent trajectory is summarized using a principled set of descriptors - including curvature, path signatures, persistent homology, and recurrent networks -that capture multiscale geometric, topological, and dynamical features. These features drive downstream prediction in a modular, interpretable, and end-to-end trainable framework.

We evaluate Neurospectrum on simulated and experimental datasets. It tracks phase synchronization in Kuramoto simulations, reconstructs visual stimuli from calcium imaging, and identifies biomarkers of obsessive-compulsive disorder in fMRI. Across tasks, Neurospectrum uncovers meaningful neural dynamics and outperforms traditional analysis methods.

## 1 Introduction

Decoding brain signals and linking them to behavior and disease states is a central challenge in cognitive neuroscience. However, the high-dimensional, complex, and spatiotemporally structured nature of neural signals—compounded by measurement noise—makes them difficult to interpret and analyze (Luo, 2021; Huk and Hart, 2019). These intricate patterns are not only prominent during active cognitive states but also persist in the resting brain, highlighting a continuous interplay of neural activity even in the absence of external stimuli (Laufs et al., 2003; Vachha et al., 2024). Furthermore, stimulus-evoked responses introduce additional complexity, as external inputs dynamically modulate these intrinsic signaling processes (Logothetis et al., 2001; Bullmore and Sporns, 2009; Sadaghiani and Kleinschmidt, 2016). Understanding these multifaceted dynamics is essential for advancing our knowledge of brain function and dysfunction.

To address this challenge, we introduce Neurospectrum - a multi-step framework that integrates geometry, topology, and signal processing to learn rich spatiotemporal representations of neural activity. Combining measures such as curvature, topological summaries, and RNNs, Neurospectrum captures meaningful differences in signal structure across conditions in both synthetic and real-world neural datasets. By incorporating differentiable components, the framework supports end-to-end training and generates attention maps that highlight brain regions critical for distinguishing experimental conditions, offering interpretable insights into neural dynamics.

Traditional methods for analyzing neural activity - such as principal component analysis (PCA), independent component analysis (ICA), and correlation metrics - often rely on simplifying assumptions like linearity, orthogonality, or independence, which rarely hold in neurobiological data (Laubach et al., 1999; Beckmann and Smith, 2004; Brendel et al., 2011; Cunningham and Yu, 2014). Correlation-based metrics (e.g., Pearson, Spearman) are widely used to infer functional connectivity (Smith et al., 2009; Honey et al., 2009), but are limited to linear or monotonic associations and assume stationarity, thus missing dynamic, nonlinear, and context-dependent neural interactions (Honey et al., 2009; Breakspear, 2017). These methods often fail to detect transient synchronization, hierarchical organization, or the flexible reconfiguration of brain networks (Bassett and Sporns, 2017; Higley, 2024). Graph-based approaches like Laplacian Eigenmaps can uncover community structures (Messaoud et al., 2025; Yang et al., 2025), but are sensitive to preprocessing and typically overlook temporal evolution.

To address dynamics, models such as HMMs characterize discrete brain states over time (Ahrends et al., 2022; He et al., 2023), while Granger causality infers directed influences (Seth et al., 2015). Yet both require strong assumptions—e.g., linearity, sufficient data, or proper model specification—and are prone to spurious inferences under noise or shared inputs. Dynamic functional connectivity approaches compute time-resolved correlations but suffer from noise sensitivity and parameter selection issues (Jiang et al., 2022). More generally, many techniques involve spatial averaging or temporal downsampling, limiting their ability to detect fine-scale patterns and reducing interpretability and robustness, particularly in noisy or heterogeneous data (Deco et al., 2011; Sadaghiani and Kleinschmidt, 2016).

Nonlinear manifold learning offers an alternative, capturing complex interactions without strong assumptions. For example, Gao et al. (2021) showed that nonlinear embeddings reveal fine-grained spatial patterns in latent brain dynamics often missed by PCA. While methods like these and those in Gonzalez-Castillo et al. (2023) capture transient synchrony and nonlinear dependencies, they typically lack explicit temporal modeling. T-PHATE addresses this by integrating diffusion-based autocorrelations into manifold learning, preserving smooth temporal trajectories and revealing large-scale state transitions (Busch et al., 2023). However, it does not explicitly model hierarchical dynamics or provide insight into spatial feature importance, limiting interpretability and multiscale analysis.

Recently, geometric and topological methods—such as wavelet transforms and persistent homology—have emerged as powerful tools for characterizing multiscale structure in brain activity. These methods are well-suited to the complexity of neural signals, capturing both fine-grained and global patterns across spatial and temporal domains. Motivated by this, we developed a framework for learning low-dimensional, interpretable representations of neural dynamics, useful for downstream applications such as identifying signaling motifs, memory retrieval, and disorder classification.

Our method, Neurospectrum, captures the spatiotemporal dynamics of neural activity through a three-stage pipeline. First, we construct a spatial graph from neural recordings and apply an attention-enhanced graph wavelet transform to extract multiscale spatial representations. Next, we learn a low-dimensional latent trajectory using a T-PHATE-regularized autoencoder, enabling joint optimization of spatial and temporal features while preserving geometric and temporal continuity. Finally, we extract complementary summaries of the latent dynamics using geometric (path signatures, curvature), topological (persistent homology, Betti curves), and temporal (RNN-based) descriptors, which are then used for downstream tasks via a feed-forward network. This modular design allows individual components to be adapted or replaced based on the application. Moreover, because several of these descriptors are inherently differentiable, Neurospectrum can be trained end-to-end, allowing the architecture to to selectively focus on the aspects of neural dynamics most salient to the task.

We evaluated Neurospectrum using mathematical models, simulations, and experimental data. First, we demonstrated that Neurospectrum representations can predict the parameters of the Kuramoto model (Kuramoto, 1975), which simulates oscillatory behavior driven by excitatory-inhibitory coupling between neurons. Next, we applied Neurospectrum to Ca^2+^ imaging data from layer 4 of the primary visual cortex in mice, showing that it can accurately reconstruct spatial frequency patterns of sinusoidal visual stimuli. Lastly, we used Neurospectrum to analyze resting-state fMRI data collected before and after a cognitive task in individuals with obsessive-compulsive disorder (OCD) and healthy controls (Ma et al., 2021). We found that Neurospectrum captured significant post-task reconfiguration in neural dynamics, particularly through increases in trajectory curvature - an effect that was more pronounced in individuals with OCD. Additionally, the model’s attention mechanism highlighted brain regions previously implicated in OCD. It also uncovered sex-specific differences in these patterns, revealing distinct regional signatures of post-task reorganization of neural activity in male and female participants.

Together, these results demonstrate the broad applicability of Neurospectrum to analyze complex spatiotemporal neural activity across a range of conditions and subject cohorts. By effectively capturing both local and global dynamics of brain activity, Neurospectrum provides a powerful framework to identify distinct neural activity motifs associated with cognitive tasks, sensory stimuli, and neurological disorders. The ability to model and classify brain activity patterns at multiple scales opens new avenues to understand the mechanisms underlying neurodevelopmental conditions, such as autism spectrum disorder, as well as other brain disorders. Furthermore, the framework’s capacity to fit mathematical models to neural signaling offers potential for developing predictive tools that can inform interventions and therapeutic strategies. Overall, Neurospectrum lays the groundwork for integrating topological and geometric methods into neuroscience, paving the way for more robust and interpretable analyses of brain dynamics in both research and clinical applications.

## 2 Results

### 2.1 Neurospectrum architecture and overview

Neurospectrum comprises three main components, each capturing a distinct aspect of the spatiotemporal structure in the data:

#### Spatial embedding

We construct a spatial graph where nodes represent brain regions, cell positions, or sensor locations, depending on the dataset. Edges are defined based on anatomical adjacency, physical proximity, or structural connectivity, while time-lapse neural signals - such as electrophysiological recordings, calcium traces, or oscillator phases - serve as node features. We then apply a graph wavelet transform, which we imbue with attention, to obtain a multi-resolution descriptor spatial representations of neural activity. The learnable attention mechanism highlights key nodes by assigning higher weights to regions with salient signal patterns across time, enhancing interpretability for different cognitive and motor tasks. The use of diffusion wavelets ensures that the representation is permutation equivariant to node ordering, making it robust to variations in sensor indexing, electrode placement, or inter-subject anatomical differences. Moreover, the multiscale nature of the wavelet transform enables the capture of hierarchical structure in neural activity, from local synchrony within micro-circuits to distributed coordination across larger networks.

#### Temporal embedding

The resulting multiscale spatial embeddings are compressed into a low-dimensional trajectory using a geometry-preserving autoencoder, capturing the temporal evolution of the system. To model the dynamics of neural activity, we learn a low-dimensional latent trajectory using a T-PHATE-regularized autoencoder. T-PHATE (Busch et al., 2023) is a manifold-aware dimensionality reduction technique that preserves both local geometric structure and temporal continuity by leveraging diffusion-based autocorrelations. We selected T-PHATE for its ability to denoise high-dimensional temporal data and reveal smooth, interpretable trajectories that reflect intrinsic neural dynamics. Rather than applying T-PHATE as a standalone preprocessing step—which would disconnect the latent trajectory from upstream spatial representations—we integrate it as a regularization term within the autoencoder’s objective. This design allows gradients to propagate through the entire pipeline, including the graph wavelet transform with attention, enabling the attention weights to be learned in a data-driven manner. As a result, the model jointly optimizes both the spatial encoding and the latent temporal trajectory, yielding representations that capture the full spatiotemporal structure of neural activity.

#### Geometrical and topological representations

We leverage topological and geometric tools to analyze the temporal dynamics in the latent trajectory. These include: (i) path signatures, which encode the geometry of trajectories through iterated integrals capturing displacement, curvature, and higher-order interactions (Chen, 1958); (ii) curvature, which reflects the smoothness and directional changes of the trajectory; (iii) Betti curves derived from persistent homology (Zomorodian and Carlsson, 2005), which summarize the birth and death of topological features such as connected components and loops across scales; and (iv) representations learned via recurrent neural networks (RNNs), which capture memory-dependent temporal dependencies in the dynamics. Each summary type provides a complementary view of neural dynamics, and they are computed independently from the latent representations. These summaries are then fed into a feed-forward architecture tailored to specific downstream tasks. This modular design not only enables the integration of multiple perspectives on brain dynamics but also supports differentiable summarization paths - such as recurrent networks or differentiable topological layers - that can be seamlessly incorporated and trained end-to-end.

Each of these components is described in detail in the following sections.

### 2.2 Spatial embedding via graph wavelet transform with attention

We begin by encoding the spatial structure of neural activity at a given timepoint *t*_*i*_. The input is a signal *X*(*t*_*i*_) defined on a graph *G* = (*V, E*), where *V* = {*v*_1_, … , *v*_*M*_ } denotes either brain regions or individual cells, depending on the dataset. The construction of the edge set *E* depends on the modality of the data: for brain activity data (e.g. fMRI), we use an anatomical adjacency graph, where each node represents a distinct brain region and edges connect spatially neighboring regions based on a predefined anatomical atlas. For calcium imaging data, we construct a cell adjacency graph, where nodes correspond to individual cells and edges are drawn between cells that share a physical boundary, as determined by their spatial segmentation.

Let *A* denote the resulting adjacency matrix and *D* the corresponding degree matrix. The signal value at node *v*_*ℓ*_ and time *t*_*i*_ is denoted by *x*(*v*_*ℓ*_, *t*_*i*_). First, we apply a learnable attention mechanism that assigns importance weights *α*_*ℓ*_ to each node, resulting in a reweighted signal:

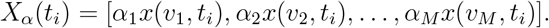

These reweighted signals are then passed through a cascade of wavelet transforms built from powers of the lazy random walk operator:

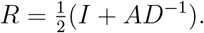

The operator *R*^*j*^ encodes the distribution of *j*-step random walks on the graph, capturing signal diffusion across increasingly large neighborhoods. We define wavelet operators at multiple scales as:

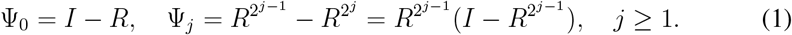

These operators isolate changes in signal across spatial scales. Inspired by techniques in graph signal processing (Gao et al., 2019; Tong et al., 2024), we apply a deep cascade of wavelet transforms interleaved with a pointwise absolute value nonlinearity:

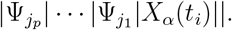

The absolute value operation is applied element-wise to the intermediate output at each step. This operation is used in place of traditional activation functions to preserve energy and oscillatory features. We now define the resulting wavelet coefficients, which serve as multiscale spatial features at time *t*_*i*_.

Zeroth-order coefficients correspond to the attention-reweighted signal:

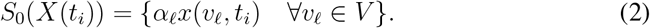

where *α*_*l*_ denotes the learnable attention on note *v*_*l*_.

First-order coefficients are obtained by applying wavelet operators to the reweighted signal:

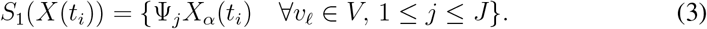

Second-order coefficients capture interactions between fluctuations at different scales:

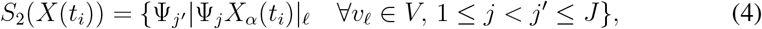

where | · |_*ℓ*_ denotes the absolute value taken at vertex *v*_*ℓ*_. These coefficients capture signal transitions at various spatial resolutions.

These capture interactions between fluctuations at different scales - e.g., how local transitions propagate to larger neighborhoods. Each of these operations is performed independently for every timepoint *t*_*i*_. The resulting embeddings for first- and second-order coefficients have shapes *M × J* and *M ×* 1*/*2*J*(*J −* 1), respectively. Finally, we concatenate all scattering orders to compute the spatial embedding:

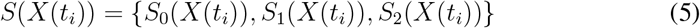

### 2.3 Temporal embedding via T-PHATE regularized autoencoder

To compress the spatial embeddings *S*(*X*(*t*_*i*_)) obtained from the graph wavelet transform, we use a geometry-preserving autoencoder regularized by T-PHATE (Busch et al., 2023). In other words, the encoder enforces that manifold distances between low dimensional embeddings of different timepoints is preserved in the latent space. This autoencoder captures the temporal evolution of multiscale spatial patterns in a low-dimensional latent space while encouraging embeddings that preserve both local geometry and global temporal structure. The architecture consists of an encoder *E* and a decoder *D*, each implemented as multilayer perceptrons (MLPs).

The encoder maps the multiscale spatial embeddings to a *d*-dimensional latent representation:

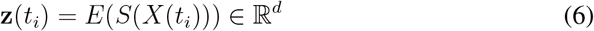

where **z**(*t*_*i*_) captures temporally meaningful, geometry-aware features at timepoint *t*_*i*_. The decoder reconstructs the original wavelet embedding from the latent space:

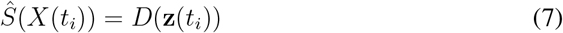

#### Reconstruction loss

The autoencoder is trained to minimize the reconstruction error between the original and decoded embeddings:

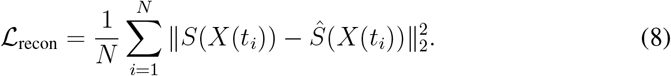

#### T-PHATE regularization

To regularize the latent space, we incorporate a loss term based on T-PHATE (Busch et al., 2023), a variant of the PHATE (Moon et al., 2019) algorithm designed to preserve both spatial and temporal structures in time-series data. T-PHATE constructs two diffusion operators:

- **P**_*D*_: captures the geometric similarities between timepoints based on the spatial patterns of the graph signals.
- **P**_*T*_ : a diffusion operator derived from the autocorrelation of the input time-series signal, capturing temporal dependencies across time lags.

A dual-view diffusion operator is then constructed by alternating spatial and temporal diffusion, **P** = **P**_*D*_**P**_*T*_ . This unified diffusion process defines pairwise diffusion distances *D*_diff_(*i, j*) between timepoints *t*_*i*_ and *t*_*j*_. The T-PHATE regularization loss encourages the Euclidean distances between latent embeddings of timepoint to match these distances:

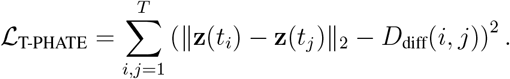

Details on the computation of the T-PHATE distances from the graph data are provided in the Computational Methods section. The total loss function combines the reconstruction and regularization terms:

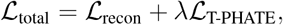

where *λ* is a tunable hyperparameter that controls the strength of the regularization.

### 2.4 Geometrical and topological summaries of latent trajectories

The latent trajectories, **z**(*t*), derived from the encoder network of our autoencoder, serve as compact representations of the multiscale spatiotemporal features extracted from the original brain signals. To interpret the dynamics encoded in these trajectories, we extract geometric, temporal, and topological features that offer complementary perspectives on their shape and evolution (Fig. 1B). Curvature captures local bending and directional changes, reflecting transitions in neural or cognitive state. Path signatures, based on iterated integrals, summarize higher-order geometric features such as displacement, loops, and enclosed areas in a coordinate-invariant manner. Topological features derived from persistent homology characterize global structure, including recurring patterns (loops) and distinct activity phases (connected components), and how these persist across scales. Recurrent neural networks are used to model temporal dependencies, capturing information about the ordering and timing of latent transitions. These descriptors are concatenated to form a unified representation, which is then passed to a feed-forward network for downstream classification or regression.

**Figure 1:**
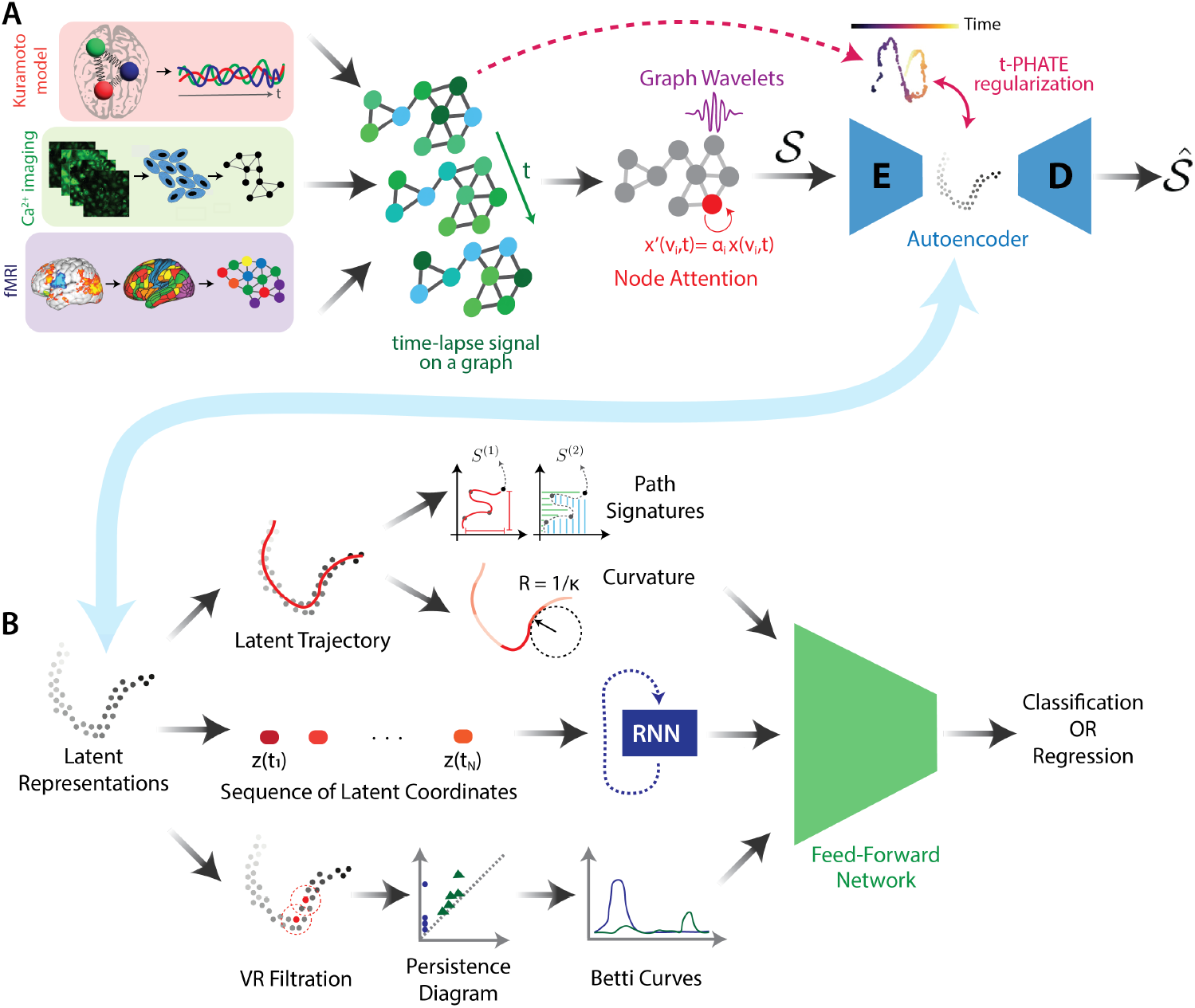
Overview of the Neurospectrum architecture. (A) We model neural dynamics from diverse sources - Kuramoto simulations, calcium imaging, or fMRI - as time-lapse signals on a graph. These signals are embedded using a deep graph wavelet transform with node attention, producing multiscale spatiotemporal features. These features are encoded using a T-PHATE- regularized autoencoder to obtain latent trajectories that preserve temporal structure. (B) From the resulting latent representations, we compute a range of geometric and topological summaries of the latent trajectory, including: path signatures (iterated integrals capturing higher-order interactions), curvature (local bending), Betti curves (topological persistence over filtration scales), and representations from recurrent neural networks (RNNs). These descriptors are concatenated and passed to a feed-forward network for downstream classification or regression tasks. The architecture is fully modular - individual summary features can be added, removed, or replaced depending on the task or data modality. Moreover, when differentiable summaries are used, the entire pipeline can be trained end-to-end, allowing the encoder, latent representations, and downstream predictors to be jointly optimized for task-specific performance.

While many of these descriptors - such as path signatures, curvature, and RNNs - are differentiable and amenable to gradient-based learning, others like Betti curves are fundamentally non-differentiable due to their discrete, combinatorial nature. Nonetheless, recent advances have introduced differentiable approximations of persistent homology, enabling the integration of topological features into end-to-end trainable models (Scoccola et al., 2024). Indeed to render Neurospectrum into an end-to-end trainable different framework the non-differentiable modalities can be excluded.

Below, we describe each of these feature representations in detail:

#### Curvature (*κ*)

To quantify local directional changes along the latent trajectory **z**(*t*), we compute curvature using a discrete circle-fitting approach (see Section 4.1.5). At each timepoint, curvature is defined as the inverse of the radius of the osculating circle that best fits a small window of points around it (Supplementary Fig. S1). This yields a localized estimate of trajectory bending, capturing transitions that may correspond to shifts in neural or cognitive states.

#### Path Signatures (PS

To capture higher-order geometric features and the temporal ordering of states in **z**(*t*), we compute path signatures - a sequence of iterated integrals that provide a coordinate-invariant representation of the trajectory’s shape (Chen, 1958). For a latent trajectory **z**(*t*) *∈* ℝ^*d*^, the *k*-th order path signature is a tensor of the form:

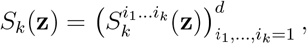

where each component

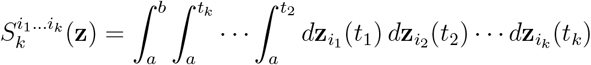

is an iterated integral along the coordinate directions *i*_1_, … , *i*_*k*_. These integrals capture progressively higher-order geometric interactions along the trajectory. First-order terms (*k* = 1) quantify net displacement along each coordinate axis, providing a coarse summary of the trajectory’s extent. Second-order terms (*k* = 2) capture the signed area swept out between pairs of coordinates, encoding aspects of curvature and directional co-dependence. Third-order terms (*k* = 3) accumulate volumetric contributions from triplets of coordinates, reflecting torsion-like effects and coordinated interactions across three dimensions. These concepts are illustrated in Supplementary Fig. S2, which shows how path signatures capture displacement, area, and volume integrals from a latent trajectory in ℝ^3^. Importantly, this formulation is invariant to reparameterizations and preserves the temporal ordering of events, making it especially well-suited for summarizing complex latent dynamics in neural time series.

#### Recurrent Neural Network (RNN)

To capture temporal dependencies within the latent trajectory, we train a recurrent neural network (RNN) on the sequence {**z**(*t*_1_), … , **z**(*t*_*n*_)}. The RNN learns internal representations that reflect the evolving dynamics of the system over time. Its final hidden state provides a compact, fixed-length summary of the trajectory, which can then be used for downstream prediction tasks. This approach is especially well-suited for settings where the order and timing of latent state transitions are critical. Architectural and training details are provided in the Computational Methods section.

#### Betti Curves (BC)

To characterize the global topological structure of the latent trajectory **z**(*t*) in a manner that is robust to small deformations and deformations, we extract features based on persistent homology (Carlsson, 2009). Given the set of latent embeddings **z**(*t*_*i*_), we treat these as a point cloud in latent space and construct a Vietoris–Rips filtration based on pairwise Euclidean distances. At each filtration scale *s*, we build a simplicial complex by including a *k*-simplex for every (*k* + 1) points whose pairwise distances are all less than or equal to *s*. A simplex is a generalization of a triangle or tetrahedron to arbitrary dimensions: a 0-simplex is a point, a 1-simplex is a line segment, a 2-simplex is a filled triangle, and so on.

As the filtration scale *s* increases, the Vietoris–Rips complex grows by incorporating higher-order simplices, forming a nested sequence of simplicial complexes that captures the multiscale topological structure of the point cloud. Persistent homology tracks how topological features, such as connected components, loops, and voids, emerge and disappear across this sequence. The birth of a feature is recorded at the smallest scale *s* where it appears in the complex, and its death is marked by the scale at which it merges with an older feature or is filled in by higher-dimensional simplices. These birth–death pairs are collected into a persistence diagram, a multiset of points (*b*_*i*_, *d*_*i*_) plotted in ℝ^2^, where each point corresponds to a topological feature and its lifespan across the filtration. Longer-lived features, which persist across a wider range of scales, are typically considered more robust or meaningful. We summarize these diagrams using Betti curves, defined as:

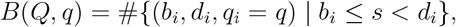

which count the number of *q*-dimensional features present at each scale *s* (Zomorodian and Carlsson, 2005). To generate a fixed-length representation, we discretize the filtration parameter and compute Betti curves in dimensions 0 and 1, which are then concatenated to form the final topological feature vector.

Together, these descriptors provide a feature-rich, multiscale summary of the geometry and topology of latent dynamics. In Neurospectrum, all extracted features are flattened, concatenated into a unified vector, and passed into a multilayer perceptron (MLP) for classification or regression. While the full model leverages this integrated representation, our ablation experiments evaluate the individual contributions of each summary by isolating them and assessing their predictive performance independently.

#### Computational Complexity

Let *n* denote the number of timepoints and *d* the dimensionality of the latent trajectory **z**(*t*). Curvature estimation is linear in the number of timepoints, with complexity 𝒪 (*n*). Computing path signatures up to order *k* scales as 𝒪 (*n* · *d*^*k*^) due to the exponential growth in feature terms. Training recurrent neural networks (RNNs) on the latent trajectory incurs a cost of 𝒪 (*n* · *d* · *h*) per sequence, where *h* is the hidden state dimension. Betti curve computation using Vietoris–Rips filtrations has worst-case exponential complexity in *d*, though practical implementations leverage sparsity and truncation strategies to remain tractable. Additional implementation details and complexity analysis can be found in the Methods.

### 2.5 Neurospectrum achieves state-of-the-art performance across multiple neural decoding tasks

We compared Neurospectrum against a broad set of baseline methods that reflect the major methodological approaches used in neural data analysis, including linear decomposition, temporal modeling, graph-based embeddings, dynamic connectivity, and topological analysis. Linear methods such as Principal Component Analysis (PCA) and Independent Component Analysis (ICA) reduce dimensionality by projecting neural activity onto directions of maximal variance or statistical independence. Temporal models, including Hidden Markov Models (HMMs) and Granger causality (GC) (Granger, 1969), infer latent state transitions or directed interactions over time, but often rely on simple parametric assumptions. Graph-based models construct correlation graphs using Pearson (Pearson, 1895) or Spearman (Spearman, 1904) coefficients and apply either spectral embeddings (GSE, using Laplacian Eigenmaps) or graph neural networks (GCNs) to extract spatial features - though these typically assume fixed connectivity and do not model temporal evolution. Dynamic methods like dynamic functional connectivity (dFC) track transient co-activation patterns over sliding windows but summarize temporal variation in a coarse, aggregated manner. Lastly, topological methods such as zig-zag persistent homology (ZZ) (Carlsson and de Silva, 2010) analyze the evolving shape of latent trajectories using overlapping temporal windows, but they do so independently of the spatial embedding. Details on these methods and their computational complexity are provided in Sections 4.3 and 4.4.

To systematically evaluate the design choices in Neurospectrum, we designed three tasks, each constructed to isolate and probe specific aspects of the latent trajectory representations described earlier. These test cases allow for targeted interrogation of the model’s ability to capture periodicity, disentangle overlapping signals, and detect transitions in coordinated dynamics - hallmarks of complex neural activity.

In the first task (Fig. 2A(i)), a single sinusoidal signal was injected at a source node and diffused across the graph with added Gaussian noise. The goal was to regress the input frequency from the resulting activity, a task requiring sensitivity to periodic structure. Neurospectrum achieved the lowest mean squared error (MSE), outperforming ICA and dFC, which performed competitively due to their sensitivity to independent and transient patterns.

**Figure 2:**
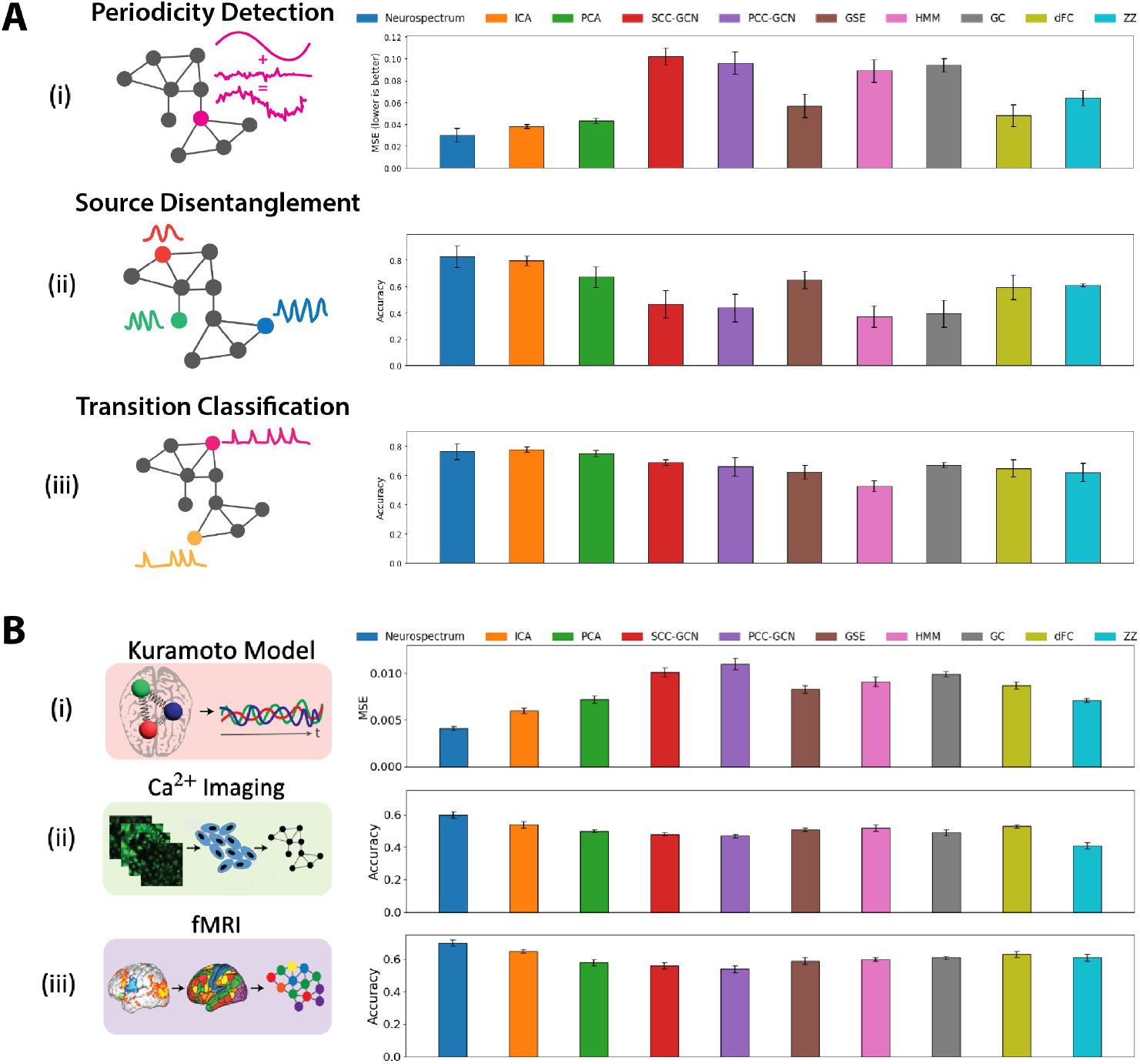
Comparison of model performance across synthetic and real-world tasks. (A) Synthetic benchmarks designed to evaluate specific capabilities of each model. (i) Periodicity Detection: A sinusoidal signal is injected at a single source node and diffused across the graph; the task is to regress its frequency. Performance is reported as mean squared error (MSE), lower is better. (ii) Source Disentanglement: Multiple sinusoidal signals of distinct frequencies are introduced at different nodes; the task is to classify the number of unique sources based on the resulting activity. Accuracy is reported; higher is better. (iii) Transition Classification: We simulate transitions from asynchronous to synchronous (collective spiking) dynamics; the task is to classify whether a transition occurred. Accuracy is reported; higher is better. (B) Real-world evaluation tasks. (i) Kuramoto Model: The task is to predict the coupling strength parameter from oscillator time series. Performance is reported as mean squared error (MSE). (ii) Ca^2+^ Imaging: The task is to classify the spatial frequency and orientation of sinusoidal grating stimuli from Calcium activity patterns in the primary visual cortex. Accuracy is reported; higher is better (iii) fMRI: The task is to classify subjects as either OCD or healthy control based on resting-state fMRI data. Accuracy is reported; higher is better.

In the second task (Fig. 2A(ii)), multiple sinusoids of distinct frequencies were introduced at different nodes. The goal was to classify the number of sources, requiring the model to disentangle overlapping spatial and temporal signals. Neurospectrum again achieved the highest accuracy, whereas graph-based methods (SCC-GCN and PCC-GCN) and temporal models (HMM, GC) struggled with overlapping dynamics.

The third task (Fig. 2A(iii)) simulated spontaneous transitions in neural dynamics - specifically, a shift from asynchronous spiking to globally synchronized activity, akin to transitions observed during neural entrainment or seizure onset. The goal was to classify whether such a transition occurred based on observed time series. Graph Spectral Embeddings (GSE), which project activity onto a static Laplacian eigenbasis, failed to capture dynamic changes in coordination that occur outside fixed spatial modes. Similarly, Hidden Markov Models (HMM), though adept at modeling discrete latent states, struggled to represent continuous, nonlinear transitions in high-dimensional space. PCA and ICA performed comparably well in this task, likely because the transition to global synchrony produces a strong low-dimensional structure, marked by increased shared variance and reduced independence across signals, that is readily captured by these linear decomposition methods. Nonetheless, Neurospectrum’s spatiotemporally regularized latent trajectories were well-suited to detecting such global shifts in dynamics, achieving the highest accuracy in this task.

Across all three settings, Neurospectrum consistently outperformed all baselines, achieving the lowest prediction error or highest classification accuracy in each case. This performance advantage stems from its integrated design: multiscale spatial encoding via graph wavelets with attention, temporally regularized latent trajectories using T-PHATE, and a rich set of geometric and topological summaries. While some baselines, such as ICA and dFC, performed reasonably well due to their sensitivity to independent or transient patterns, they lacked the inductive bias needed to capture spatial structure or continuous temporal evolution. Graph-based models struggled with static connectivity assumptions, and temporal or topological methods fell short due to their limited expressiveness.

To further dissect the contributions of individual components, we performed a series of ablation studies on the same tasks (see Section 4.5 and Fig. S3). These ablations validate the necessity of each module in our architecture, showing marked drops in performance when either the spatial encoding, temporal regularization, or trajectory summarization is removed or simplified.

Building on these controlled evaluations, we next tested Neurospectrum on more complex tasks involving nonlinear simulations and experimental recordings. In the Kuramoto oscillator task (Fig. 2B(i)), we compared models tasked with inferring the coupling strength parameter from collective phase dynamics, assessing their ability to capture emergent global patterns. In the calcium imaging task (Fig. 2B(ii)), models decoded the spatial frequency and orientation of visual stimuli, requiring sensitivity to fine-scale, stimulus-evoked activity in the mouse visual cortex. Finally, in the resting-state fMRI classification task (Fig. 2B(iii)), the goal was to distinguish OCD patients from healthy controls based on resting-state brain activity, emphasizing the need to extract neural signatures from noisy and task-free brain dynamics. Neurospectrum outperformed all baselines across these diverse settings, demonstrating strong generalization to real-world neural data. Further details are provided in Sections 2.6, 2.7, and 2.8.

### 2.6 Neurospectrum quantifies synchrony in coupled oscillators

We first validated Neurospectrum on the Kuramoto model (Kuramoto, 1975), a classical framework for studying collective synchronization in networks of coupled oscillators. Originally developed to model phase synchronization in physical and chemical systems, the Kuramoto model has since been widely used to understand neuronal populations and the emergence of coherent brain oscillations such as alpha and theta waves (Strogatz, 2000; Acebrón et al., 2005; Zhang et al., 2018). These oscillatory patterns are believed to play a critical role in cognitive function, attention, and memory, motivating our choice of the Kuramoto model as a canonical benchmark for evaluating spatiotemporal structure in neural-like dynamics. The dynamics of the system are governed by:

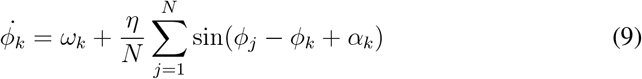

where *ϕ*_*k*_ is the phase of oscillator *k, ω*_*k*_ *∼* 𝒩 (1, *β*) its intrinsic frequency, *α*_*k*_ a phase shift, and *η* the coupling strength. At low *η*, oscillators evolve independently. At higher *η*, coupling induces synchrony, and for sufficiently high values, the population becomes globally entrained, with all oscillators locking into a common phase. To study these transitions, we simulated a network of 100 oscillators arranged on a hexagonal grid, evolving their dynamics over 2000 timepoints with a timestep of Δ*t* = 0.1, for a total simulated duration of *t* = 200.

We applied Neurospectrum to the resulting time series to extract latent spatiotemporal representations of network activity across varying levels of coupling. We found that Neurospectrum outperforms baseline methods in recovering the underlying coupling parameter *η* from observed dynamics (Section 4.6). To contextualize these results, we examined the raw behavior of the system at representative coupling strengths (Fig. 3A). At *η* = 0.1, oscillators remain desynchronized, resulting in a broad phase distribution. At *η* = 0.5, clusters of locally synchronized oscillators emerge. At *η* = 0.9, global synchronization dominates, with all oscillators aligning in phase.

**Figure 3:**
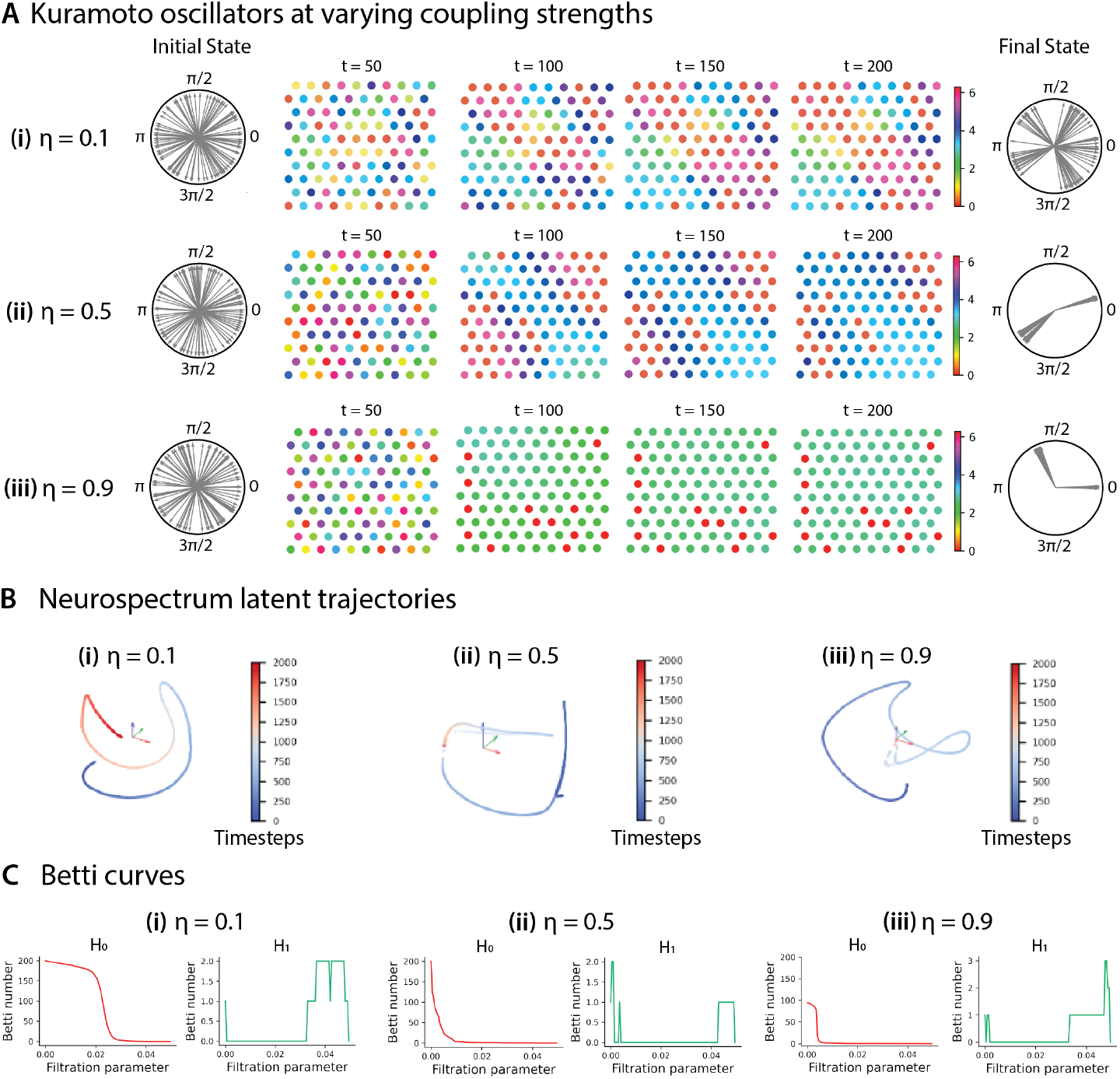
Neurospectrum quantifies synchrony and entrainment in simulations of weakly coupled Kuramoto oscillators. (A) Snapshots from simulations of 100 Kuramoto oscillators arranged in a hexagonal grid at three different coupling strengths *η* = 0.1, 0.5, 0.9, where each oscillator interacts with up to 6 neighbors. The phase of each oscillator (ranging from 0 to 2*π*) is color-coded. Polar plots on the left and right show the initial and final phase distributions, respectively. At low coupling (*η* = 0.1, i), oscillators evolve independently, maintaining diverse phases. At intermediate coupling (*η* = 0.5, ii), subsets of oscillators begin to synchronize, forming phase-aligned clusters. At high coupling (*η* = 0.9, iii), global entrainment emerges as all oscillators converge to a shared phase. (B) Neurospectrum-derived latent trajectories illustrate the system’s evolution over time, colored by timestep. For low coupling, the trajectory progresses smoothly, reflecting gradual, unsynchronized changes. At intermediate coupling, the trajectory converges onto a localized region of latent space, signifying partial synchronization. For strong coupling, the trajectory quickly collapses into a single point, indicating rapid and stable population-level entrainment. (C) Betti curves computed from the latent trajectories quantify topological features across scales. Dimension-0 homology (*H*_0_, red) reflects component connectivity, while dimension 1 (*H*_1_, green) captures loop structures. At higher coupling strengths, the steep decline in *H*_0_ indicates tightly clustered embeddings, consistent with shared oscillator state and synchronization.

Latent trajectories extracted by Neurospectrum reflect these transitions in state space (Fig. 3B). For low coupling, the trajectory smoothly progresses through latent space, with each timestep embedded adjacent to its neighbors - indicating a system undergoing continuous, unsynchronized evolution. At intermediate coupling, the trajectory gradually collapses onto a compact region, signifying partial synchrony. At high coupling, the trajectory rapidly converges to a single point, where it remains stationary, capturing the emergence of a stable, shared state across the population.

Topological summaries of the latent trajectories further characterize these dynamics. In Fig. 3C, we show Betti curves derived from a Vietoris-Rips filtration applied to the latent trajectories, where pairwise Euclidean distances between timepoints serve as the metric for simplicial complex construction. As the filtration parameter increases, points (i.e., timepoints in the latent space) that are close begin to merge into connected components, and larger-scale topological features such as loops eventually emerge.

The dimension-0 homology group (*H*_0_) characterizes the number of connected components at each scale, and its rank, the 0-th Betti number, quantifies how many such components exist at a given filtration value. For a compact latent trajectory - such as one where the system quickly converges to a shared state - timepoints are close together, and components merge early during filtration, resulting in a low and rapidly declining *H*_0_ Betti curve. In contrast, for a dispersed trajectory that evolves continuously without convergence, components remain separated over a longer range of filtration values, resulting in high Betti numbers that persist. Consistent with this, we observe that at low coupling (*η* = 0.1), the latent trajectory remains extended, with many disconnected components that merge only at higher filtration scales - reflected in a slow decline in *H*_0_ Betti curve (Fig. 3C(i)). At intermediate (*η* = 0.5) and high (*η* = 0.9) coupling strengths, the *H*_0_ Betti curve declines much more steeply (Fig. 3C(ii)-(iii)), indicating that the latent trajectory is highly clustered, consistent with convergence to a narrow region or single point in latent space as the oscillators synchronize.

The dimension-1 homology group (*H*_1_) captures 1D holes or loops in the latent trajectory, with the 1st Betti number quantifying the number of such independent cycles. Across all coupling regimes, we observe nontrivial *H*_1_ features that emerge at higher filtration values, reflecting the arc-like structure of the trajectory in latent space. These loops are indicative of turns or cycles in the latent trajectory that arise during transitions in system state, such as the formation of new clusters of synchronous oscillators or the merging of previously separate groups. The presence and persistence of these *H*_1_ features highlight structured dynamics even in the absence of full synchrony.

To further evaluate the capability of Neurospectrum to detect transitions in synchrony, we conducted an additional experiment in which the coupling strength *η* was dynamically modulated over time (Fig. S4). In this setting, *η* was set to zero initially, increased to one during two brief intervals to induce transient synchrony, and then reset back to zero. This design mimics real-world neural dynamics where periods of coordinated activity are interspersed with desynchronized fluctuations. The latent trajectory extracted by Neurospectrum captures these transitions: during high *η*, the trajectory converges to a compact region of latent space associated with synchrony, while periods of desynchrony trace out distinct paths that eventually return to this region upon resynchronization (Fig. S4B). The RNN summary trained to regress *η* from segments of the latent trajectory effectively separates these regimes, with synchronized timepoints forming distinct clusters in the learned embedding (Fig. S4C). These findings underscore the RNN’s capacity to encode temporally extended dependencies that correspond to underlying changes in network-level coupling, and demonstrate the broader utility of Neurospectrum in tracking dynamic, reversible shifts in collective behavior.

Taken together, these results show that Neurospectrum provides a compact and informative representation of network synchrony across a range of coupling regimes, and can successfully recover global parameters that govern emergent collective behavior. Importantly, inferring the coupling strength *η* from observed dynamics constitutes an inverse problem: rather than simulating dynamics from known parameters, we aim to recover a hidden global variable from noisy, locally interacting, graph-structured time series. This task is particularly challenging due to the many-to-one, nonlinear nature of the forward map, the stochasticity introduced by initial phases and natural frequencies, and the delayed or subtle emergence of synchrony - especially at intermediate coupling. Neurospectrum’s ability to solve this problem highlights its strength in capturing latent structure in complex, nonlinear dynamical systems.

### 2.7 Neurospectrum reconstructs sinusoidal patterns shown to mice from visual cortex recordings

Next, we evaluated Neurospectrum on publicly available two-photon calcium imaging data from the mouse visual cortex (Garner, 2014) to assess its ability to decode structured visual stimuli from neural activity. In this experiment, head-fixed mice with cranial windows were shown sinusoidal grating patterns that systematically varied in either spatial frequency (distance between bars) or orientation (angle of the bars). Each stimulus was presented for 3 seconds, followed by a 2-second inter-stimulus interval, and repeated 8 times per condition. Calcium imaging data were acquired from layer 4 of the primary visual cortex at approximately 30 Hz.

The grating patterns consisted of parallel stripes whose brightness varied according to a sinusoidal function (Fig. 4A), a widely used class of stimuli in vision research due to their ability to elicit reliable and tunable neural responses.

**Figure 4:**
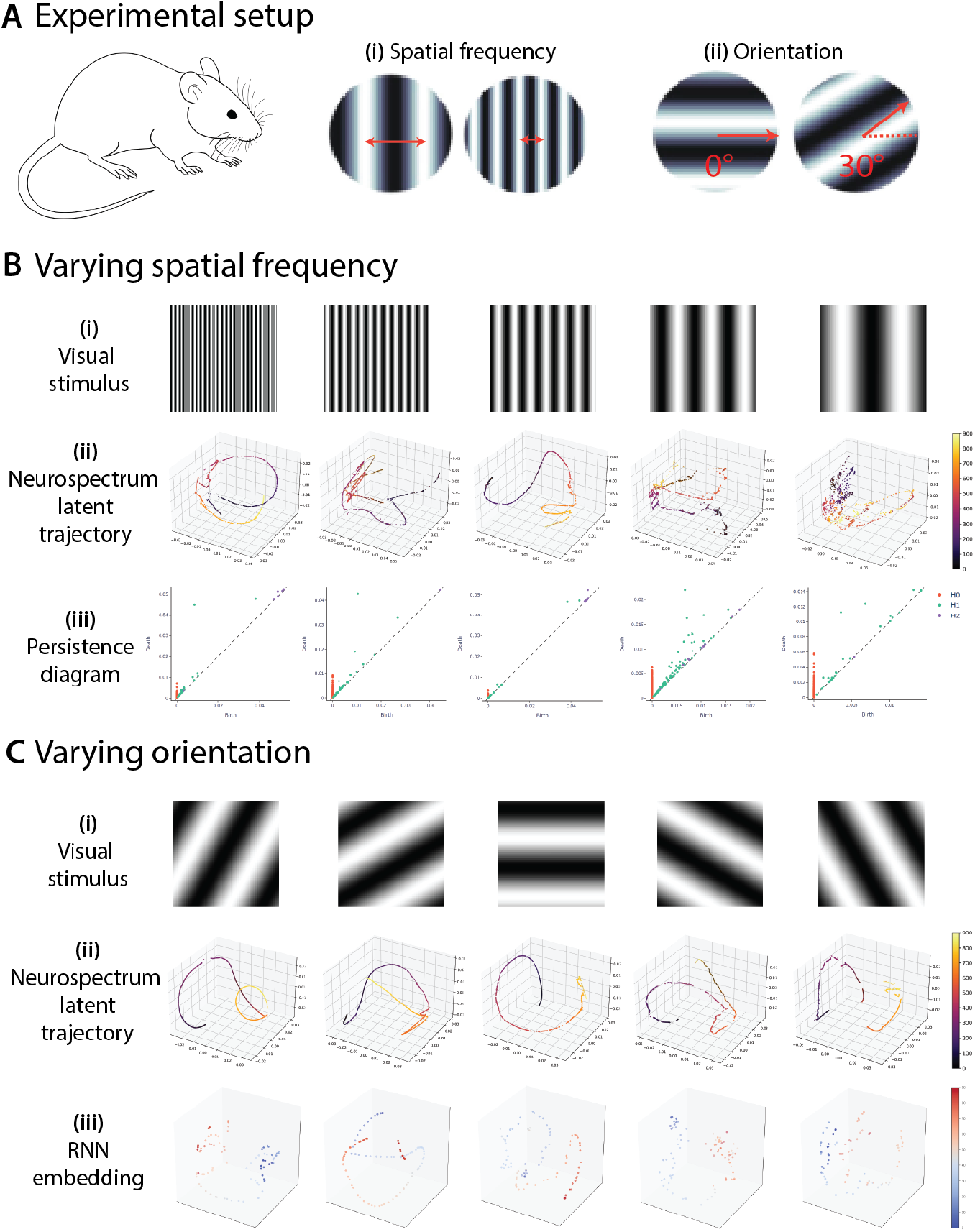
Signatures of neural activity in layer 4 of the primary visual cortex in response to sinusoidal visual stimuli. (A) Experimental setup for sinusoidal grating patterns of various (i) spatial frequencies and (ii) orientations, displayed on a computer screen to head-fixed mice fitted with head-plates and a cranial window. (B) Grating patterns with (i) varying spatial frequencies, (ii) their Neurospectrum latent trajectories, and (iii) the persistent diagram of the latent trajectories. Low-frequency stimuli exhibited more prominent off-diagonal components in both dimension-0 (*H*_0_) and dimension-1 (*H*_1_) homology groups, indicating greater topological complexity. (C) Grating patterns with (i) varying orientations, (ii) their Neurospectrum latent trajectories, and (iii) the RNN embedding of the latent trajectories. Neurospectrum produced topologically similar embeddings, further RNN embeddings showed noisier transitions for gratings with orientations greater than 90^°^ (clockwise from vertical), compared to those below.

To assess how well Neurospectrum captures stimulus-induced dynamics, we trained it to learn low-dimensional latent trajectories from the high-dimensional neural recordings. Across both spatial frequency and orientation conditions, Neurospectrum consistently outperformed baseline models in predicting stimulus identity from brain activity (Section 4.7).

We first examined responses to gratings of varying spatial frequency (Fig. 4B(i)). The geometry of the learned latent trajectories (Fig. 4B(ii)) was strongly modulated by frequency content. High-frequency stimuli produced smooth, low-dimensional curves with coherent temporal evolution, indicative of synchronized and structured neural dynamics. In contrast, low-frequency stimuli resulted in noisier, more disorganized trajectories with no well-defined curvature.

To quantify these differences, we computed persistent homology on the latent trajectories and summarized their topological structure using Betti curves (computed from the persistence diagrams shown in Fig. 4B(iii)). High-frequency stimuli exhibited strong 1-dimensional topological features (e.g., prominent loops in *H*_1_), while low-frequency stimuli showed a proliferation of short-lived features in both *H*_0_ and *H*_1_, indicative of fragmented or transient activity patterns.

We next examined neural responses to gratings with varying orientations (Fig. 4C(i)). While the topology of the latent trajectories remained relatively consistent across orientations (Fig. 4C(ii)), differences in temporal structure were captured by the RNN-based embeddings (Fig. 4C(iii)). Notably, orientations greater than 90^°^ (measured clockwise from vertical) produced more dispersed and temporally heterogeneous embeddings, suggesting greater variability in stimulus-driven dynamics.

Together, these results highlight the modular strength of our framework. Betti curves capture global geometric and topological properties of neural trajectories, while recurrent embeddings extract fine-grained temporal structure. By combining these complementary representations, Neurospectrum effectively distinguishes between stimulus features that manifest through different aspects of neural activity - spatial frequency through geometry and topology, and orientation through temporal variability - offering a powerful and interpretable approach for understanding how stimuli are encoded at population-level neural activity.

### 2.8 Neurospectrum identifies task-induced changes in brain dynamics and classifies OCD with high accuracy

To investigate how task performance affects intrinsic brain dynamics, we applied Neurospectrum to resting-state fMRI data collected before and after a novel task designed to probe both perceptual and value-based decision-making processes (PVDM) (Ma et al., 2021). In this dataset, participants made decisions using the same individually calibrated visual stimuli under two conditions: perceptual discrimination and value-guided choice. The dataset includes both healthy control (HC) participants and unmedicated individuals diagnosed with obsessive-compulsive disorder (OCD). Previous analyses of this dataset revealed gender-specific alterations in decision formation in OCD, particularly among males, who showed increased caution and reduced evidence accumulation efficiency compared to matched controls. A graph of brain regions was constructed using the DiFuMo atlas (Dadi et al., 2020), with each node representing a cortical or subcortical region and edges connecting spatially adjacent regions (Fig. S5A). At each timepoint, the BOLD signal was averaged within each region to define a signal over the graph. This graph-structured signal was transformed into multiscale spatial embeddings using a graph wavelet transform with attention (Fig. S5B), as described earlier. These embeddings were then encoded into a low-dimensional latent trajectory via a T-PHATE-regularized autoencoder that preserved both spatial structure and temporal continuity (Fig. S5C). From these trajectories, we extracted a suite of geometrical, topological, and recurrent features, including curvature (Fig. S5D) and Betti curves (Fig. S5E), which were used for downstream analysis. As detailed in Section 4.8, Neurospectrum outperforms all baseline methods in classifying OCD versus HC individuals based on post-task resting-state activity. We now focus on results obtained using Neurospectrum to characterize group and sex-specific alterations in intrinsic brain dynamics.

We first examined whether the latent trajectories captured by Neurospectrum reflected task-induced changes in brain dynamics. Comparing pre and post-task trajectories revealed marked differences in curvature: specifically, post-task trajectories exhibited more frequent and localized sharp turns, corresponding to elevated curvature values (Fig. 5A). These changes were robustly detected by curvature (*κ*) and path signatures (PS), both of which are sensitive to transient, localized features in the dynamics.

**Figure 5:**
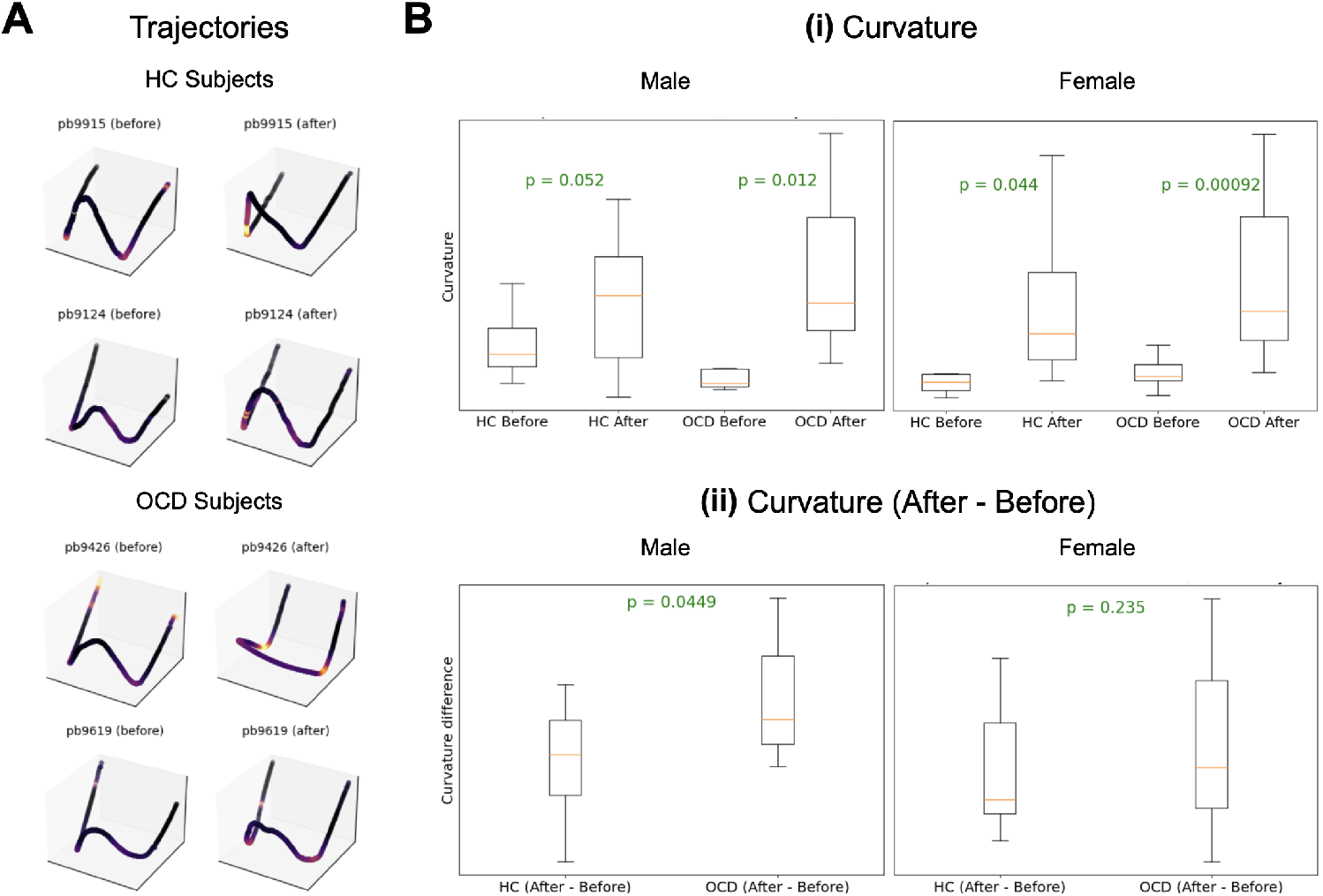
(A) Neurospectrum trajectories for healthy control (HC) and OCD subjects from fMRI dataset. (B) (i) Box plots of curvature computed from Neurospectrum trajectories for HC and OCD subjects, before and after the task, shown separately for males and females; (ii) Box plots of curvature increases after the task, with a more pronounced effect in OCD subjects of both sexes. *p*-values are computed using the Mann–Whitney U test, with lower values observed in male subjects.

The observed increase in curvature was significantly greater in OCD participants than in HCs across both sexes (Fig. 5B(i)), suggesting that individuals with OCD may exhibit heightened dynamical reconfiguration or instability in response to cognitive engagement. This effect was especially pronounced in males, where the separation between OCD and HC groups based on curvature differences was larger than in females, pointing to a possible sex-specific signature. This finding is consistent with prior work (Ma et al., 2021) highlighting sex-dependent neural alterations in OCD.

Further stratification by sex revealed that although both males and females exhibited post-task curvature increases, the effect was significantly more pronounced in males overall (Fig. 5B(ii)), suggesting stronger or more heterogeneous patterns of reconfiguration in the latent dynamics of male brains. These sex differences also manifested in downstream classification: using curvature features alone, OCD vs. HC classification yielded a statistically significant *p*-value of 0.0449 in males, but not in females (*p* = 0.235), suggesting that the discriminative power of curvature-based features may differ by sex.

To further improve discriminative power and interpretability, we leveraged the differentiable nature of the RNN module - among other components in our framework - to integrate classification directly into the model and enable end-to-end training. This design enables the learnable attention mechanism to highlight brain regions that contribute most to the differences between OCD and HC participants. By jointly optimizing reconstruction and classification objectives, the model learns spatiotemporal representations that are not only faithful to the multiscale brain dynamics but also maximally discriminative. This end-to-end framework allows the attention map to reflect diagnostically relevant brain regions, providing interpretable insights into region-specific alterations associated with OCD. This highlights the use of Neurospectrum as a fully differentiably neural network architecture.

Finally, we examined the attention weights learned by the spatial encoder, which highlight brain regions most influential in differentiating OCD from HC participants. These attention maps revealed a subset of cortical and subcortical regions - such as the Putamen Posterior in female participants - that are implicated in the altered brain dynamics associated with OCD (Fig. 6A), consistent with findings from Kubota et al. (2016). In male participants, high attention weights in the Superior Frontal Sulcus suggest this region plays a key role in the heightened separability between HC and OCD groups, in line with recent findings by Doyle et al. (2024). Furthermore, we observed marked sex differences in the spatial distribution of attention weights. Notably, the Middle Frontal Gyrus Anterior and the Putamen exhibited the most prominent divergence between male and female participants, suggesting potential sex-specific patterns of functional engagement in OCD (Fig. 6B). These findings underscore the importance of incorporating sex as a biological variable in the study of neural dynamics and highlight distinct spatial mechanisms that may underlie post-task reconfiguration in OCD across sexes.

**Figure 6:**
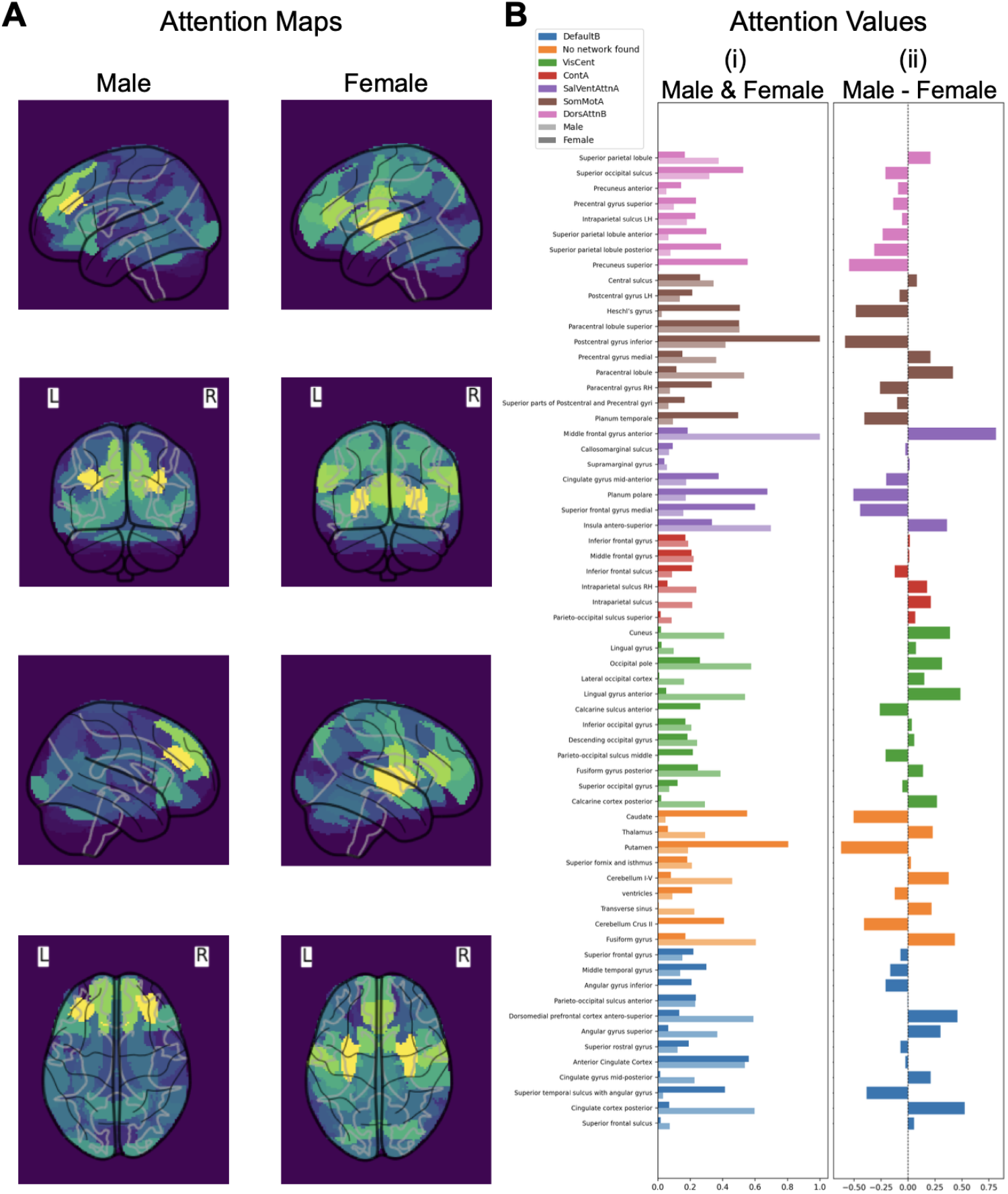
(A) Cortical mesh visualization of learned attention values across brain regions, using parcels from the DiFuMo atlas, shown separately for male and female subjects. Attention values concentrates in Superior Frontal Sulcus in male subjects and Putamen Posterior in female subjects. (B) (i) Histogram of attention values across brain regions for males and females and (ii) their differences. Middle Frontal Gyrus Anterior and the Putamen exhibited the most prominent divergence between male and female subjects.

## 3 Discussion

We introduce Neurospectrum a geometric and topological deep learning framework for analyzing complex spatiotemporally patterned neural dynamics through learned latent trajectories. The framework begins by modeling the measurements substrate (consisting of voxels or other measurement units) as an anatomical near-neighbors graph and applying a library of graph wavelet transforms with an attention mechanism that learns an importance score wavelets centered at each node in the brain graph. While we use anatomical adjacency in this work, the underlying graph could also be constructed from functional connectivity, structural connectomics, or other domain-specific measures of inter-region relationships, depending on the data modality and task. The wavelet transforms capture both local and global patterns of neural activity across multiple spatial scales. These multiscale representations are then compressed into low-dimensional time trajectories using a regularized autoencoder that preserves the manifold structure of the data and respects its temporal continuity via a regularization that enforce the embedding to respect a neural manifold learning method called T-PHATE. Effectively this creates an extensible encoder that can map new points into the T-PHATE space. From these latent trajectories, we compute a suite of geometric and topological descriptors - including path signatures, curvature, Betti curves, and recurrent neural network representations - each capturing distinct structural or dynamical features of neural activity over time that we finally concatenate to feed to prediction-head MLP networks.

A key insight from our work is that different trajectory descriptors capture different aspects of the underlying dynamics and therefore excel at different tasks. For instance, path signatures and Betti curves proved most effective for capturing periodicity and cyclic structure in oscillator simulations, while curvature and RNN-based features were more sensitive to transient events such as sudden transitions from random to coordinated neural activity. This complementarity highlights the strength of a modular design that can flexibly adapt to the demands of diverse neural phenomena, rather than relying on a one-size-fits-all representation. Crucially, the set of trajectory descriptors can be customized or extended to suit specific applications. If the descriptors are differentiable, such as RNN encoders or parameterized geometric functions, the entire framework becomes end-to-end differentiable, allowing the model to be trained jointly with task-specific objectives for optimal performance.

Neurospectrum was systematically evaluated on three representative datasets, each designed to probe distinct aspects of neural dynamics. In our first study, we examined whether Neurospectrum could recover the coupling strength of coupled oscillators simulated using the Kuramoto model, a widely used dynamical system for studying synchronization in neuronal populations. This setting presents a challenging inverse problem, as small changes in an unobserved global coupling parameter can lead to highly nonlinear and emergent network-wide synchronization patterns. Neurospectrum accurately inferred this coupling strength from the observed local phase trajectories, demonstrating its ability to extract global dynamical properties from local, noisy measurements. In calcium imaging data from mouse visual cortex, Neurospectrum decoded properties of visual stimuli - such as spatial frequency and orientation - based on population-level neural activity, showing that its latent representations preserve stimulus-relevant structure in the dynamics. Finally, in resting-state fMRI data collected before and after a cognitive task, Neurospectrum identified task-induced reconfiguration in brain dynamics and uncovered sex-specific markers of OCD, with path signatures and curvature capturing subtle but reliable differences in trajectory shape between healthy and clinical populations. Together, these results demonstrate that Neurospectrum generalizes across both synthetic and real neural systems, and across both continuous regression and discrete classification tasks.

Importantly, Neurospectrum does not merely produce high-dimensional embeddings - it yields interpretable, task-relevant signatures of neural dynamics. The temporal encoder’s attention mechanism highlights informative brain regions, and the geometric/topological summaries capture properties such as trajectory complexity, reconfiguration, and recurrence. These interpretable features enabled accurate prediction of synchrony in oscillator networks, reconstruction of visual stimuli from calcium imaging, and classification of individuals with OCD from resting-state fMRI.

By integrating multiscale spatial filtering with unsupervised temporal modeling and interpretable latent trajectory analysis, Neurospectrum offers a general and extensible framework for decoding complex neural activity. The model’s ability to operate across simulation, calcium imaging, and fMRI data - and its flexibility in supporting diverse downstream tasks - suggests broad applicability to problems in systems neuroscience, cognitive science, and psychiatric research. Beyond the brain, the core ideas underlying Neurospectrum may also prove useful for analyzing other biological systems where information is encoded in dynamic patterns over structured spatial domains, such as signaling networks, collective cell migration, or ecological dynamics.

## 4 Methods

### 4.1 Mathematical Background and Implementation Details

#### 4.1.1 Manifold Learning and Diffusion Geometry

A useful assumption in representation learning is that high dimensional data originates from an intrinsic low dimensional manifold that is mapped via nonlinear functions to observable high dimensional measurements; this is commonly referred to as the manifold assumption. Formally, let ℳ^*d*^ be a hidden *d*-dimensional manifold that is only observable via a collection of *n ≫ d* nonlinear functions *f*_1_, … , *f*_*n*_ : ℳ^*d*^ *→* ℝ that enable its immersion in a high dimensional ambient space as *F* (ℳ^*d*^) = {**f** (*z*) = (*f*_1_(*z*), … , *f*_*n*_(*z*))^*T*^ : *z ∈* ℳ^*d*^} ⊆ ℝ^*n*^ from which data is collected. Conversely, given a dataset *X* = {*x*_1_, … , *x*_*N*_ } *⊂* ℝ^*n*^ of high dimensional observations, manifold learning methods assume data points originate from a sampling 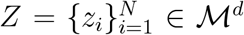 of the underlying manifold via *x*_*i*_ = **f** (*z*_*i*_), *i* = 1, … , *n*, and aim to learn a low dimensional intrinsic representation that approximates the manifold geometry of ℳ^*d*^.

To learn a manifold geometry from collected data, scientists often use the diffusion maps construction of (Coifman and Maggioni, 2006) that uses diffusion coordinates to provide a natural global coordinate system derived from eigenfunctions of the heat kernel, or equivalently the Laplace-Beltrami operator, over manifold geometries. This construction starts by considering local similarities defined via a kernel 𝒦 (*x, y*), *x, y ∈ F* (ℳ^*d*^), that captures local neighborhoods in the data. We note that a popular choice for 𝒦 is the Gaussian kernel exp(*−*∥*x − y*∥^2^*/σ*), where *σ >* 0 is interpreted as a user-configurable neighborhood size. However, such neighborhoods encode sampling density information together with local geometric information. To construct a diffusion geometry that is robust to sampling density variations, we use an anisotropic kernel:

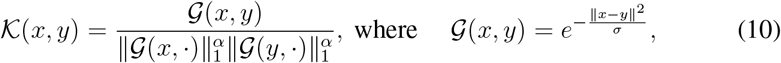

as proposed in Coifman and Maggioni (2006), where 0 *≤ α ≤* 1 controls the separation of geometry from density, with *α* = 0 yielding the classic Gaussian kernel, and *α* = 1 completely removing density and providing a geometric equivalent to uniform sampling of the underlying manifold. Next, the similarities encoded by 𝒦 are row-normalized to define transition probability matrix **P**_*ij*_ = *p*(*x*_*i*_, *x*_*j*_) where

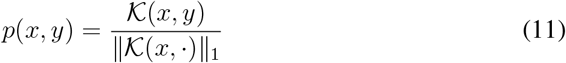

This matrix **P** describes a Markovian diffusion process over the intrinsic geometry of the data.

#### 4.1.2 T-PHATE Regularization

The timelapse signal *X* on a graph forms a matrix of dimensions *T ×M* , where *T* is the number of time points and *M* is the size of the signal at each timepoint. To visualize the signal over time, we calculate a distance matrix *D*(*i, j*) = ∥*X*(*t*_*i*_) *− X*(*t*_*j*_)∥_2_ using the Euclidean distance between timepoint signals. *D* is converted from a distance matrix into a probability matrix *P* using Eq. (10) and Eq. (11) for the Markovian random-walk diffusion process. The diffusion timescale, *t*_*D*_, is computed, which specifies the number of steps taken in the random-walk process. This parameter provides a tradeoff between encoding local and global information in the embedding, where a larger *t*_*D*_ corresponds to more steps than a smaller *t*_*D*_. *t*_*D*_ is computed automatically using the spectral or von Neumann entropy of the diffusion operator. Next, we compute the T-PHATE distance between the distributions in the *i*-th and *j*-th rows of 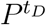, the *t*_*D*_-step random walk over *P* :

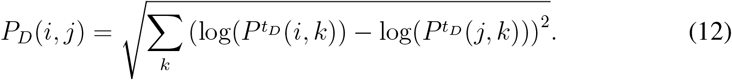

Next, we compute the autocovariance of the *T × M* timelapse signal using *T −* 1 lags, resulting in a *M ×* (*T −* 1) matrix. Next, we average across the timelapse signal *M* to obtain a single vector *c* of autocorrelation at each lag. We then construct the temporal affinity matrix *A*, by identifying the lag *l*_max_ where *c* = 0:

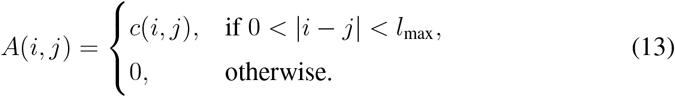

We convert the autocorrelation matrix into the transition probability matrix *P*_*T*_ by row-normalizing the affinity matrix *A* and powering it to *t*^*D*^. Finally, we alternate the spatial and temporal diffusion operators to construct a dual-view diffusion operator *P* = *P*_*D*_*P*_*T*_ . We embed *P* using metric MDS to obtain 3-D T-PHATE coordinates, *E*(*t*) = (*E*_1_(*X*(*t*)), *E*_2_(*X*(*t*)), *E*_3_(*X*(*t*))) for each time point *t* for visualization, which can also be viewed as the point cloud *E* = {*E*(*t*_1_), *E*(*t*_2_), … , *E*(*t*_*n*_)} for subsequent topological data analysis.

#### 4.1.3 Persistent homology and topological data analysis

Topological data analysis (TDA) refers to techniques for understanding complex datasets by their topological features, i.e., their connectivity (Carlsson, 2009). Here we focus on the topological features of a data graph where the simplest set of topological features are given by the number of connected components *b*_0_ and the number of cycles *b*_1_, respectively. Such counts, also known as the Betti numbers, are coarse graph descriptors that are invariant under graph isomorphisms. Their expressivity is increased by considering a function *f* : *V →* ℝ on the vertices of a graph, *G* = (*V, E*), with vertex set *V* and edge set *E*. Since *V* has finite cardinality, so does the image im*f* , i.e., im*f* = {*w*_1_, *w*_2_, … , *w*_*n*_}.

Without loss of generality, we assume that *w*_1_ *≤* · · · *≤ w*_*n*_. We write *G*_*i*_ for the subgraph induced by filtering according to *w*_*i*_, such that the edges satisfy 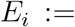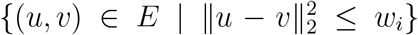 . The subgraphs *G*_*i*_ satisfy a nesting property, as *G*_1_ ⊆ *G*_2_ ⊆ · · · ⊆ *G*_*n*_. When analyzing a point cloud, the vertices of each *G*_*i*_ arise from spatial coordinates for the data and *w*_*i*_ constitutes a distance threshold between points, such that *G*_*n*_ is a fully-connected graph, containing all the vertices from *V* . This is commonly known as the Vietoris-Rips (VR) filtration.

It is then possible to calculate topological features alongside this filtration of graphs, tracking their appearance and disappearance as the graph grows. If a topological feature is created in *G*_*i*_, but destroyed in *G*_*j*_ (it might be destroyed because two connected components merge, for instance), we represent this by storing the point (*w*_*i*_, *w*_*j*_) in the persistence diagram 𝒟_*f*_ associated to *G*. Another simple descriptor is given by the Betti curve of dimension *d* of a diagram 𝒟, which refers to the sequence of Betti numbers of dimension *d* in 𝒟, evaluated for each threshold *w*_*i*_.

#### 4.1.4 Recurrent Neural Network (RNN)

To model temporal dependencies within the latent representations, we employ a Recurrent Neural Network (RNN) to process sequential data. Given an input sequence of latent representations *z*(*t*_1_), *z*(*t*_2_), … , *z*(*t*_*n*_), the RNN extracts temporal features that encode the underlying dynamics of the sequence. Unlike autoencoder-based approaches that aim to reconstruct future latent states, our model utilizes the RNN purely as a feature extractor, where the final hidden state serves as a learned representation of the entire sequence.

The extracted representation is then fed directly into a feedforward neural network (FFN) for classification. This architecture allows the model to leverage the sequential structure of the latent space while maintaining efficiency in downstream tasks. Formally, given a sequence of latent states *Z* = [*z*(*t*_1_), *z*(*t*_2_), … , *z*(*t*_*n*_)], the RNN transforms it into a fixed-dimensional representation *h*_*n*_, which is subsequently passed to the classifier:

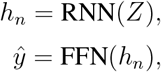

where 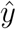 denotes the predicted class label. The RNN and FFN are trained jointly to optimize the classification objective, ensuring that the learned temporal representations are directly useful for distinguishing between different classes. This approach enables effective utilization of sequential dependencies without explicitly reconstructing future latent states.

#### 4.1.5 Computation of curvature

Curvature is a measure of how much the curve deviates from a straight line. In other words, the curvature of a curve at a point is a measure of how much the change in a curve at a point is changing, meaning the curvature is the magnitude of the second derivative of the curve at given point. A plane curve given by Cartesian parametric equations *x* = *x*(*t*) and *y* = *y*(*t*), the curvature kappa, sometimes also called the “first curvature” (Kreyszig, 1991), is defined by

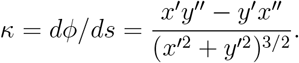

Here we consider curves in 3-dimensional Euclidean space, specified parametrically by *x* = *r* cos *t* and *y* = *r* sin *t*, which is tangent to the curve at a given point. The curvature is then

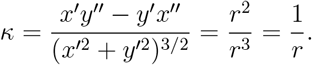

For curvature at point *p*, we fit a 1-dimensional circle *S*^1^(*p, r*) centered at *p* with radius *r* in the plane spanned by principal components of the T-PHATE trajectory. The inverse of radius 1*/r* gives the curvature at *p* (see Fig. S1).

To elaborate, at each point *p* of the curve *C*, we select a local neighborhood of points around *p*. The size of this neighborhood, a user-defined hyper-parameter (set here to 5% of the total curve length), determines the number of points sampled symmetrically around *p*. The neighborhood is then centered by subtracting the mean of these points from each point, ensuring that the analysis is performed relative to the center of mass. Next, Singular Value Decomposition (SVD) is applied to the centered neighborhood, yielding two vectors that span the local plane and a normal vector perpendicular to this plane. A circle is then fitted to the points in the local plane using a least-squares method. The curvature at *p* is subsequently calculated as the reciprocal of the radius (1*/r*) of the fitted circle, assuming that locally the trajectory approximates a circular arc. This procedure is repeated for all points along the trajectory, giving a curvature profile across the entire curve.

#### 4.1.6 Graph construction for calcium imaging data

We represent the imaged tissue as a graph *G* = {*V, E*}, consisting of nodes *v*_*i*_ *∈ V* and edges (*v*_*j*_, *v*_*k*_) *∈ E*, where each node *v*_*i*_ represents a cell and a pair of nodes *v*_*j*_ and *v*_*k*_ is connected with an edge based on a predefined criterion. For Ca^2+^ signaling data from the primary visual cortex of mice, we connect nodes that are spatially adjacent (within 2 *µm* of each other), as the flow of signals is thought to be between spatially proximal cells. The connectivity of graph *G* can be described by its adjacency matrix **A**, where **A**_*ij*_ = 1 if *v*_*i*_ and *v*_*j*_ are connected and 0 otherwise. The degree of each vertex is defined as a diagonal matrix D, where 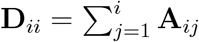.

Graph signals can associate with each node or edge in a graph. In the Ca^2+^ signaling data, the signals associated with cell *v*_*i*_ is the normalized Ca^2+^ fluorescence intensity at each timestep *t*. Since every cell has a related Ca^2+^ signal, this signal *X*(*v*_*i*_, *t*) is defined over the whole graph for timestep *t*.

### 4.2 Computational Complexity of Neurospectrum

Below we describe the computational complexity of all steps of Neurospectrum.

At each timepoint *t*_*i*_, the spatial signal *X*(*t*_*i*_) is reweighted by a learned attention vector ***α*** *∈* ℝ^*M*^ , and passed through a cascade of graph wavelet transforms constructed using powers of the lazy random walk operator 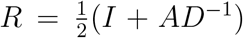. The transform extracts multiscale features by applying dyadic powers of *R*, as defined in Eq. (1), at zeroth-, first-, and second-order levels. For a graph with *M* nodes and *J* dyadic scales, the per-timepoint cost of computing first- and second-order transforms is 𝒪 (*MJ* + *MJ*(*J −* 1)*/*2) = 𝒪 (*MJ*^2^). Across *T* timepoints, the total cost scales as 𝒪 (*TMJ*^2^). These matrix-vector operations are efficiently implemented using sparse matrix routines (e.g., in PyTorch or SciPy), and parallelized across both timepoints and nodes.

To compress the time-indexed spatial embeddings *S*(*X*(*t*_*i*_)) into a *d*-dimensional latent trajectory **z**(*t*_*i*_), we use an autoencoder comprising multilayer perceptrons (MLPs) with hidden dimension *h*. The encoder and decoder each require 𝒪 (*dh*^2^) operations per timepoint, resulting in a total cost of 𝒪 (*Tdh*^2^) for a forward-backward pass over *T* timepoints. The additional regularization term aligns latent distances with T-PHATE distances, requiring the computation of a dual-view diffusion operator and pairwise distances, which scales as 𝒪 (*T* ^2^*d*) but can be precomputed and stored.

The computation of geometrical and topological representations introduces several post-hoc steps, which can all be run in parallel. Curvature estimation scales linearly with *T* , while path signatures up to order *k* scale as 𝒪 (*Td*^*k*^), though in practice *k ≤* 3 is sufficient for expressivity. Persistent homology of the latent trajectory (for Betti curves) is computed using Vietoris–Rips complexes, with worst-case complexity 𝒪 (*T* ^3^) but efficient implementations, e.g., Ripser (Tralie et al., 2018), GUDHI (Maria, 2025), make computation feasible for moderate *T* (typically under 500). The final feed-forward classifier operates on the concatenated feature vector and has negligible cost relative to earlier stages.

While Neurospectrum introduces higher computational overhead than simpler baselines, this is offset by substantial gains in representational power, as demonstrated in our empirical evaluations.

### 4.3 Baseline methods for comparison

To evaluate the ability of Neurospectrum to capture meaningful spatiotemporal dynamics of neural activity, we compared its performance against a suite of baseline methods that are widely used in the analysis of neural signals. These baselines span classical linear techniques, graph-based spatial embeddings, temporal modeling, and methods that estimate time-varying or directed connectivity. All baseline methods produce fixed-length feature vectors, which were used as input to a multilayer perceptron (MLP) for downstream classification or regression tasks.

#### Principal Component Analysis (PCA

PCA was applied over the spatial domain, reducing the dimensionality of each timepoint’s brain activity vector by projecting it onto the top principal components that collectively explained at least 90% of the total variance. The resulting low-dimensional vectors were concatenated across time to yield a spatiotemporal feature representation for each sample. PCA serves as a linear baseline that captures dominant variance directions, albeit without preserving statistical independence or spatial locality.

#### Independent Component Analysis (ICA)

Similar to PCA, ICA was applied across the spatial domain, but instead decomposes the data into statistically independent sources. We used the FastICA algorithm (Hyvarinen, 1999) to extract spatially independent components and their corresponding timecourses. Each component groups together neural activity from regions that share the same temporal response patterns, thus intrinsically capturing functional connectivity. These component timecourses were concatenated across time to form a fixed-length spatiotemporal feature vector. Unlike PCA, which captures directions of maximal variance, ICA emphasizes higher-order statistical independence and is widely used in neuroimaging to identify functionally distinct neural sources (Jung et al., 2001; Calhoun et al., 2009; Robinson and Schöpf, 2013).

#### Static Correlation-Based Graphs with GCNs

To assess whether simple graph neural network architectures can extract informative structure from static functional connectivity for predictive modeling, we constructed graphs using two common measures of statistical association: Pearson correlation (PCC) (Pearson, 1895) and Spearman rank correlation (SCC) (Spearman, 1904). We computed pairwise correlations between the time series recorded at all spatial locations, resulting in a fully connected weighted graph where edge weights reflected the strength of linear or monotonic dependence.

We evaluated two graph neural network baselines, *SCC-GCN* and *PCC-GCN*, in which the raw time series at each spatial location served as the node features, and the correlation matrices defined the edge weights of the input graphs. A graph convolutional network (GCN) was used to extract node representations from these correlation-weighted graphs, with global average pooling applied to the final node embeddings to obtain a fixed-length graph-level representation. These baselines assess the predictive capability of a simple GNN architecture that relies on static correlation structure for spatial modeling.

#### Graph Laplacian Eigenmaps

We constructed a graph Laplacian *L* = *D − A* from a spatial adjacency matrix based on anatomical proximity or physical overlap, consistent with the graph used in Neurospectrum. The bottom *d* nontrivial eigenvectors of *L* were computed to form a Laplacian eigenbasis (Belkin and Niyogi, 2003), which captures local manifold structure. At each timepoint *t*_*i*_, the brain activity vector *X*(*t*_*i*_) was projected onto this eigenbasis, resulting in a *d*-dimensional embedding that encodes spatial structure. These projections were concatenated across time to produce a fixed-length spatiotemporal feature vector.

#### Hidden Markov Models (HMM)

HMMs provide a probabilistic framework for modeling temporal sequences as transitions between discrete latent states, each characterized by distinct statistical properties of the observed signal. In neural time series analysis, HMMs have been used to segment activity into recurring states that reflect coarse temporal structure or shifts in network-wide coordination (Quinn et al., 2018). These models provide a discretized view of underlying dynamics, often reducing rich spatiotemporal patterns to a limited set of state transitions.

We trained a Gaussian-emission HMM on the multivariate time series, allowing both spatial and temporal dynamics to be summarized through the model’s latent state sequence. The number of hidden states was selected using the Bayesian Information Criterion (BIC). We extracted a set of features from the fitted model, including the most probable state sequence, state transition probabilities, and fractional occupancy of each state over time. These features were concatenated to form a fixed-length vector, which was then used for downstream prediction. This baseline enables comparison to a widely-used class of temporal models that can detect discrete transitions in multivariate dynamics.

#### Granger Causality (GC)

We estimated directed functional interactions using Granger causality (Granger, 1969). A vector autoregressive (VAR) model was fit to the multivariate time series, and pairwise GC values were computed to quantify how much past activity in one region predicted future activity in another. The resulting directed adjacency matrix was vectorized and used as the feature representation. This method emphasizes directional dependencies, capturing causality-like interactions.

#### Dynamic Functional Connectivity (dFC)

To capture temporal variations in connectivity patterns within neural activity data, we employed a dynamic functional connectivity (dFC) approach based on sliding-window correlations. This approach has been widely used in neuroscience to reveal transient connectivity states and dynamic reconfigurations of brain networks that are not captured by static measures (Hutchison et al., 2013). Specifically, we computed pairwise Pearson correlations across all channels within a sliding temporal window, yielding a sequence of time-resolved connectivity matrices. From these, we extracted summary statistics describing the distribution of connectivity strengths over time, i.e. the proportion of values falling into high negative, moderate negative, low/uncorrelated, moderate positive, and high positive ranges. These features were concatenated into a fixed-length vector for downstream prediction tasks.

#### Zig-zag Persistent Homology (ZZ)

To capture how the topological structure of the Neurospectrum latent trajectory **z**(*t*) evolves over time, we use zig-zag persistent homology, a generalization of classical persistent homology that relaxes the requirement for nested filtrations (Carlsson and de Silva, 2010). Unlike standard persistence, which requires that each successive subset of data contains the previous one, zig-zag persistence can handle more flexible sequences where data can both enter and leave over time.

We begin by dividing the latent trajectory **z**(*t*_*i*_) into overlapping temporal windows. Each window forms a point cloud in latent space, from which we construct a Vietoris–Rips complex based on pairwise Euclidean distances. For instance, we define simplicial complexes over consecutive windows as:

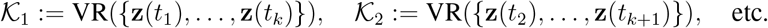

To link these temporally adjacent windows and enable tracking of features across them, we construct intermediate complexes on the union of adjacent windows:

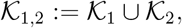

which yields a zig-zag sequence of the form:

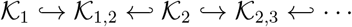

Here, arrows indicate whether points are added (forward inclusion ↪) or removed (backward inclusion ↩), allowing us to track topological features, such as connected components and loops, that persist across overlapping windows, even as the underlying data changes. The resulting zig-zag persistence diagrams summarize the birth and death of topological features across the sequence. To convert these diagrams into fixed-length numerical features, we first map each persistence point to a discretized 2D plane and apply Gaussian smoothing to obtain persistence images (Adams et al., 2016). These images, computed for dimensions 0 and 1, are then flattened and concatenated to produce a vectorized representation of the trajectory’s evolving topological structure. This method is particularly valuable in cases where the latent trajectory undergoes abrupt changes, reconfigurations, or reversals over time.

### 4.4 Computational complexity of baseline methods

Principal Component Analysis (PCA) requires computing the *M×M* covariance matrix followed by eigendecomposition. This results in a total complexity of 𝒪 (*M* ^2^*T* + *M* ^3^), where *M* is the number of brain regions and *T* is the number of timepoints. These operations are highly optimized and parallelizable via standard linear algebra libraries (e.g., NumPy, SciPy, scikit-learn, CuPy). Independent Component Analysis (ICA) builds on PCA by performing additional iterative optimization to maximize statistical independence among components. The whitening step has similar cost to PCA, while the optimization phase introduces an additional cost of 𝒪 (*IM* ^2^) for *I* iterations. Like PCA, ICA is easily parallelized across components and supports GPU acceleration in some implementations (e.g., FastICA (Hyvärinen and Oja, 2000), PyTorch).

Static correlation methods (Spearman and Pearson) compute pairwise correlations across all region pairs, with a total cost of 𝒪 (*M* ^2^*T* ). These computations are trivially parallelizable, with each correlation computed independently. Training GCN models on fully-connected graphs with edge weights obtained from SCC/PCC nvolves neighborhood aggregation and matrix multiplications in each layer. For *L* layers and node feature dimensionality *d*, the cost is 𝒪 (*LMd*^2^). We used a 3-layer GCN implemented using PyTorch Geometric (Fey and Lenssen, 2019) and DGL (Wang et al., 2019), which support batch processing and parallel execution across data samples and nodes.

Laplacian Eigenmaps project spatial patterns onto the intrinsic coordinates of the graph by computing the bottom *k* nontrivial eigenvectors of the Laplacian matrix *L* = *D − A*. The cost of eigendecomposition is up to 𝒪 (*M* ^3^) in the dense case, though this is typically reduced to 𝒪 (*kM* ^2^) using sparse solvers.

Hidden Markov Models (HMMs) model temporal dynamics through latent state sequences and require *O*(*TK*^2^) per sample, where *K* is the number of hidden states. The forward-backward algorithm is the main computational bottleneck and is parallelizable across data samples and states.

Granger Causality (GC) estimates directed relationships using vector autoregressive (VAR) models. For a VAR of order *p*, the cost scales as 𝒪 (*M* ^2^*Tp*), with one model fit per node pair. Since each VAR model is independent, this step is fully parallelizable.

Dynamic Functional Connectivity (dFC) estimates time-varying interactions by computing pairwise correlations between brain regions within overlapping temporal windows. Each window contains a fixed number of timepoints (determined by the window size *W* ) and this size is constant with respect to the overall data dimensions and does not affect the asymptotic complexity. For a dataset with *M* regions and *T* timepoints, the number of windows scales linearly with *T* , and within each window, pairwise correlations are computed for all 𝒪 (*M* ^2^) region pairs. Therefore, the total computational complexity is 𝒪 (*M* ^2^*T* ). These correlation computations are parallelized across both region pairs and time windows.

Zig-zag (ZZ) persistence involves building simplicial complexes over overlapping temporal windows and computing persistence diagrams across a sequence of inclusions. While the computation is more complex than standard persistence, it can be efficiently parallelized across time windows and is supported by optimized libraries such as Dionysus and GUDHI. Although the worst-case computational complexity is 𝒪 (*m*^3^) in the number of simplices *m*, this is significantly reduced in practice by limiting the analysis to local temporal windows and leveraging advanced matrix reduction techniques.

### 4.5 Ablation study validates modular architecture of Neurospectrum

To evaluate the individual contributions of core architectural components in Neurospectrum, we performed a series of targeted ablation experiments. Each experiment isolates a key module to assess its necessity for achieving high performance across decoding tasks. Specifically, we ablated (i) temporal regularization via T-PHATE during training, (ii) spatial encoding via the graph wavelet transform, and (iii) the use of learned latent trajectories.

We first assessed the role of temporal regularization. In the full model, T-PHATE is incorporated as a training loss that encourages the latent trajectory **z**(*t*) to preserve local temporal continuity and geometry. To ablate this component, we removed the temporal loss and instead generated low-dimensional embeddings using PCA or t-SNE applied post hoc to the encoder outputs. These variants, *NeuroSPECTRUM-PCA* and *NeuroSPECTRUM-tSNE*, consistently underperformed compared to the temporally regularized model, highlighting the benefit of incorporating temporal priors directly into training rather than relying on unsupervised dimensionality reduction after the fact.

Next, we tested the importance of spatial encoding by bypassing the graph wavelet transform and applying T-PHATE directly to the raw graph signals over time. This ablation, labeled *T-PHATE*, retained temporal structure but lacked spatially informed, multiscale features. While it achieved moderate performance, it consistently fell short of the full model, suggesting that spatial priors are essential for capturing structure in neural dynamics - particularly in settings involving localized or distributed activity patterns.

To evaluate whether a learned latent space is necessary at all, we included a purely topological baseline, *GraphPH-CROCKER*, which computes persistent homology directly on the time-varying graph signals. In this ablation, we perform a sublevel set filtration on the scalar values of each timepoint’s signal across graph nodes, yielding a sequence of persistence diagrams. These are assembled over time into a CROCKER matrix (Topaz et al., 2015), which summarizes the birth and death of topological features across time and scale. This approach avoids any encoding or compression and instead aggregates topological features using the CROCKER representation. While this baseline preserves a rich topological summary, it lacks the ability to align or smooth dynamics across time, leading to consistently lower performance compared to Neurospectrum.

In addition to architectural ablations, we evaluated the performance of each individual summary feature - Betti curves (BC), curvature (*κ*), path signatures (PS), and recurrent neural networks (RNN) - used independently in place of the full ensemble (“All”). These features capture distinct aspects of latent dynamics, and their effectiveness varies by task.

Figure S3A shows performance across the three synthetic benchmarks previously introduced in Section 2.5. The full model outperformed all ablated variants across all tasks.

In the periodicity detection task (Fig. S3A(i)), Betti curves and path signatures achieved the lowest error, as they effectively capture loop-like latent trajectories and global cyclicity - hallmarks of periodic structure. In contrast, curvature and RNN-based representations, which emphasize local dynamics or short-term memory, performed worse due to their limited ability to encode repeating structure. In the source disentanglement task (Fig. S3A(ii)), RNN features proved most effective, as they captured the evolving interaction and separation of signal sources over time. Static geometric or topological features struggled to resolve overlapping signals without an explicit temporal model. In the transition classification task (Fig. S3A(iii)), curvature-based features performed best, capturing abrupt changes in trajectory shape due to transitions into synchronized activity. RNN features also performed strongly, detecting temporal buildup preceding the transition. Although path signatures and Betti curves detected overall shifts in geometry, they were less sensitive to short, transient events.

Interestingly, in some cases, individual summary features exceeded the performance of the full ensemble (“All”), suggesting that summary selection could be task-specific. This reinforces the modular nature of Neurospectrum: different combinations of geometric and topological descriptors can be selectively applied depending on the task and data modality.

The performance gap is particularly striking when compared to the T-PHATE model, which lacks wavelet-based multiscale spatial encoding: despite capturing temporal diffusion geometry, it fails to disentangle overlapping spatial sources or detect coordinated dynamics as effectively. This suggests that wavelet-based spatial priors are essential for capturing localized neural signal propagation. The GraphPH-CROCKER baseline, which lacks both spatial encoding and latent compression, performed the worst, reinforcing the value of combining spatial filtering with temporally regularized latent representation.

To evaluate how these architectural components generalize beyond controlled synthetic settings, we applied the same ablation analysis to real-world tasks involving non-linear oscillator dynamics (Kuramoto model), stimulus-evoked neural activity (Ca^2+^ imaging), and resting-state brain recordings (fMRI). As shown in Fig. S3B, the performance trends observed in synthetic benchmarks largely hold across these domains: ablated models consistently underperform compared to the full architecture, and the relative utility of individual summary features remains task-dependent. Full results and interpretations for each of these datasets are presented in the following sections.

### 4.6 Neurospectrum outperforms baselines in recovering Kuramoto coupling

We evaluated the ability of Neurospectrum to recover the coupling strength *η* from time series data generated by the Kuramoto model. This inverse modeling task is challenging due to nonlinear dynamics, stochastic variability across replicates, and the gradual or delayed onset of synchrony, particularly at intermediate coupling strengths.

To benchmark performance, we simulated oscillator dynamics at 10 evenly spaced values of *η ∈* [0, 1], with 10 independent replicates per value. From each simulation, we extracted spatiotemporal features using Neurospectrum and trained a regression model to predict *η*, using 5-fold cross-validation.

Neurospectrum achieved the best performance across all methods (Table S1 and Fig. 2B(i)), with a mean squared error (MSE) of 0.0041 *±* 0.0002. The next-best method, independent component analysis (ICA), achieved an MSE of 0.0060 *±* 0.0003, highlighting its strength in capturing independent sources of spatially distributed activity. Principal component analysis (PCA), while effective in reducing dimensionality, underperformed (0.0072 *±* 0.0004) due to its emphasis on variance rather than functional segregation. Graph-based baselines using static correlation matrices (PCC-GCN, SCC-GCN), spectral embeddings, temporal models (HMM, Granger causality), and dynamic connectivity features (dFC) all showed higher MSEs. Zig-zag persistent homology (ZZ), which captures evolving topological structure in latent space, performed better than most baselines but still fell short of Neurospectrum and ICA.

To assess the contributions of individual components within Neurospectrum, we conducted an ablation study in which each feature - Betti curves (BC), curvature (*κ*), path signatures (PS), and recurrent neural networks (RNN) - was used in isolation (Table S2 and Fig. S3B(i)). All ablated variants underperformed relative to the full model, confirming that these features provide complementary views of the dynamics. Among single-feature models, path signatures and Betti curves performed best, underscoring their ability to capture temporal evolution and topological structure, respectively. Curvature alone was less informative, while the RNN-only model captured temporal dependencies but lacked spatial and topological context. Notably, all single-feature variants outperformed traditional baselines, emphasizing the value of the latent trajectory learned by Neurospectrum.

We also evaluated versions of Neurospectrum in which the latent trajectory was replaced by PCA, t-SNE, or T-PHATE embeddings. Each resulted in worse performance, further underscoring the importance of the multiscale, graph-structured, and temporally regularized latent space constructed by Neurospectrum. Among these alternatives, T-PHATE performed best, consistent with its use as a regularization in our model.

Taken together, these results demonstrate that Neurospectrum provides a more accurate and expressive representation for recovering global coupling strength from graph-structured oscillator dynamics than a wide range of commonly used baseline methods.

### 4.7 Neurospectrum outperforms baselines in classifying visual stimuli from neural activity

We evaluated Neurospectrum on a supervised classification task: predicting the spatial frequency and orientation of sinusoidal grating stimuli from neural activity recorded via two-photon calcium imaging. The dataset consisted of trials where drifting sinusoidal gratings were presented at 1 Hz temporal frequency and 80% contrast. Each trial was labeled with one of five spatial frequencies (0.02 to 0.32 Hz) and one of eight orientations (0° to 315° in 45° steps).

For each trial, Neurospectrum was used to extract a latent spatiotemporal trajectory from the neural recordings. A 5-layer feedforward neural network was then trained to predict stimulus identity based on this representation. As shown in Table S1 and Fig. 2B(ii), Neurospectrum achieved the highest accuracy of 0.60*±*0.02, outperforming all baseline methods. ICA was the strongest baseline (0.54 *±* 0.02), followed closely by dynamic functional connectivity (0.53 *±* 0.01) and Hidden Markov Model (0.52 *±* 0.02). These methods likely perform well due to their ability to capture distinct temporal patterns and transitions in neural activity. PCA, graph-based and topological baselines resulted in lower accuracy.

To assess the contribution of individual components within Neurospectrum, we conducted an ablation study in which each summary feature - Betti curves (BC), curvature (*κ*), path signatures (PS), and recurrent neural networks (RNN) - was used in isolation (Table S2 and Fig. S3B(ii)). Among single-feature models, Betti curves achieved the highest accuracy (0.62 *±* 0.01), slightly exceeding the full model, followed by path signatures (0.59 *±* 0.01). Curvature and RNN alone performed worse, but all ablated variants still outperformed traditional dimensionality reduction and connectivity-based baselines.

We also evaluated modified versions of Neurospectrum in which the latent trajectory was replaced by PCA, t-SNE, or T-PHATE embeddings. All replacements resulted in reduced performance, highlighting the importance of the graph-structured and temporally regularized latent space learned by Neurospectrum.

These results validate that Neurospectrum not only compresses high-dimensional calcium imaging data into informative low-dimensional trajectories but also learns representations that capture both the spatial organization (via geometric and topological features) and temporal evolution (via recurrent embeddings) of neural responses. The modular nature of the framework allows different feature types to contribute complementary information, enabling accurate decoding of complex, structured stimuli from brain activity, compared to conventional dimensionality reduction or sequence modeling approaches.

### 4.8 Neurospectrum outperforms baselines in classifying healthy vs. OCD subjects from resting-state fMRI

We assessed the ability of Neurospectrum to classify individuals with obsessive-compulsive disorder (OCD) versus healthy controls (HC) using resting-state fMRI data collected after participants completed a decision-making task. This binary classification task is particularly challenging due to the subtle, spatially distributed, and temporally unconstrained nature of resting-state brain dynamics.

Despite these challenges, Neurospectrum achieved the highest classification accuracy (0.70 *±* 0.02), outperforming all baseline methods (Table S1, Fig. 2B(iii)). Among the baselines, ICA (0.65*±*0.01) and dynamic functional connectivity (0.63*±*0.02) were the second and third best-performing methods, reflecting their sensitivity to fine-grained and temporally evolving activity patterns. However, they lack the ability to jointly model multiscale spatial structure and temporal evolution, as done by Neurospectrum’s learned latent trajectories.

Ablation analysis revealed that path signatures alone surpassed the full model (0.72*±* 0.01 vs. 0.70 *±* 0.02), indicating their effectiveness in capturing subtle trajectory deformations and transient patterns (Table S2, Fig. S3B(iii)). Notably, curvature (0.63*±*0.01) and RNN-based summaries (0.63 *±* 0.02) also outperformed most baselines. These results complement findings in Section 2.8, where differences between pre-task and posttask curvature showed significant group differences between OCD and HC subjects. Together, they suggest that local geometric changes and recurrent temporal motifs in the latent space carry discriminative signatures of OCD.

Substituting Neurospectrum’s learned latent space with conventional embeddings such as PCA, t-SNE, and T-PHATE led to reduced performance. PCA and t-SNE, in particular, yielded substantially lower accuracy (0.56 *±* 0.02 and 0.44 *±* 0.03, respectively), indicating their inability to preserve relevant spatiotemporal structure. T-PHATE performed better (0.62 *±* 0.02), likely due to its use of temporal autocorrelation to preserve continuity in the embedding. Nevertheless, none of these alternatives matched the performance of the learned latent trajectory in Neurospectrum.

Altogether, these results show that Neurospectrum is highly effective at capturing spatiotemporal brain dynamics as measured by fMRI. Its integration of topological, geometric, and recurrent features into a unified representation enables state-of-the-art performance is classifying healthy and OCD subjects and holds promise for developing data-driven biomarkers of neuropsychiatric conditions.

### 4.9 Experimental Methods

#### 4.9.1 Experimental setup for the calcium imaging dataset

We analyzed two-photon calcium imaging data originally published by Garner (2014) and made available through the CRCNS data repository. Imaging was performed using a custom two-photon microscope with resonant and galvo scanning mirrors, and excitation was provided by a femtosecond pulsed laser (MaiTai, Spectra-Physics) tuned to 910 nm. Images were acquired at *≈* 30 Hz, with each frame composed of 512 scan lines, targeting layer 4 neurons roughly 300 microns beneath the cortical surface in adult mice.

To prepare for imaging, mice were injected with a GCaMP6s calcium indicator virus (AAV1.Syn.Flex.GCaMP6s.WPRE) at postnatal day 36 and later underwent surgery to implant a head plate and a cranial window over the visual cortex. The window consisted of a glass coverslip assembly that was secured to the skull. After recovery, animals were gradually habituated to head-fixation and the imaging environment over a three-week period.

During experiments, mice were head-fixed but allowed to move on a freely spinning disk while visual stimuli were displayed to the eye opposite the recorded hemisphere. Stimuli were presented using custom Python software and consisted of full-field drifting gratings at 80% contrast. The stimuli spanned 5 spatial and 5 temporal frequencies, each shown at 8 evenly spaced orientations (0° to 315°, in 45° increments). Each unique combination was presented eight times for 3 seconds each, with 2-second inter-stimulus intervals. The monitor displaying the stimuli subtended 130° horizontally and 60° vertically and was positioned 22 cm from the mouse.

#### 4.9.2 Experimental setup for fMRI dataset

We analyzed functional MRI data from 58 adult participants who completed a novel decision-making task designed to probe both perceptual and value-based decision-making processes in parallel (the Perceptual and Value-based Decision-Making task, PVDM) using individually calibrated visual stimuli (Ma et al., 2021). The sample included 27 unmedicated individuals diagnosed with obsessive-compulsive disorder (OCD; 16 female) and 31 healthy controls (15 female), matched for age, gender, and IQ. Prior analyses of this dataset found that males with OCD, in particular, showed more cautious decision-making and less efficient evidence accumulation compared to matched control participants.

fMRI data were preprocessed using the minimal preprocessing pipeline developed by the Human Connectome Project (HCP) (Glasser et al., 2013). Subsequent processing involved segmentation of cortical and subcortical structures, alignment to the MNI-152 template (Fonov et al., 2011). Movement scrubbing and regression of nuisance variables were performed to ensure data quality.

## Data Availability

The in vivo calcium imaging dataset from layer 4 cells in the mouse primary visual cortex is available from the CRCNS portal at http://dx.doi.org/10.6080/K0C8276G. fMRI data from healthy controls and subjects with OCD are available upon request.

## Code Availability

Code for reproducing all the results presented in this paper is available on GitHub at https://github.com/KrishnaswamyLab/Neuro-SPECTRUM.

## Acknowledgments

This research was funded by Kavli Institute for Neuroscience Postdoctoral Fellowship from Yale University [D.B.], CIFAR AI Chair [Y.Z., G.W.], NIH NIGMSR01GM135929 [G.W., S.K.], NIH R01GM130847 [S.K], NSF CAREER award IIS-2047856 [S.K.], NSF DMS grant 2327211 [S.K., G.W.], NSF CISE grant 2403317 [S.K.], CRM-Simons visiting professor award [S.K.], NSERC Discovery grant 03267 [G.W.], FRQNT grant 343567 [G.W.].

## Supplementary Information

**Table S1:** Model performance and baseline comparison across three tasks: predicting coupling strength in the Kuramoto model (MSE, lower is better), decoding frequency and relative orientation of sinusoidal gratings from neural activity in primary visual cortex (PVC) (accuracy, higher is better), and classifying healthy vs. OCD individuals using resting-state fMRI data (accuracy, higher is better). The best performing model is **bolded** and the second-best is <u>underlined</u>. Mean and standard deviation computed from 5-fold cross-validation.

| Model | Kuramoto (MSE) | PVC (Acc.) | fMRI (Acc.) |
| --- | --- | --- | --- |
| **Neurospectrum** | **0.0041 ± 0.0002** | **0.60 ± 0.02** | **0.70 ± 0.02** |
| ICA | <u>0.0060 ± 0.0003</u> | <u>0.54 ± 0.02</u> | <u>0.65 ± 0.01</u> |
| PCA | 0.0072 ± 0.0004 | 0.50 ± 0.01 | 0.58 ± 0.02 |
| SCC-GCN | 0.0101 ± 0.0005 | 0.48 ± 0.01 | 0.56 ± 0.02 |
| PCC-GCN | 0.0110 ± 0.0006 | 0.47 ± 0.01 | 0.54 ± 0.02 |
| Graph Spectral Embedding (GSE) | 0.0083 ± 0.0004 | 0.51 ± 0.01 | 0.59 ± 0.02 |
| Hidden Markov Model (HMM) | 0.0091 ± 0.0005 | 0.52 ± 0.02 | 0.60 ± 0.01 |
| Granger Causality (GC) | 0.0099 ± 0.0003 | 0.49 ± 0.02 | 0.61 ± 0.01 |
| Dynamic Functional Connectivity (dFC) | 0.0087 ± 0.0004 | 0.53 ± 0.01 | 0.63 ± 0.02 |
| Zig-zag (ZZ) Persistent Homology | 0.0071 ± 0.0002 | 0.41 ± 0.02 | 0.61 ± 0.02 |

**Table S2:** Performance comparison of model ablations across three tasks: predicting the coupling strength in the Kuramoto model (MSE, lower is better), predicting the frequency and relative orientation of a sinusoidal grating pattern (accuracy, higher is better), and classifying healthy vs. OCD subjects (accuracy, higher is better). The best-performing model is shown in bold, and the second-best is <u>underlined</u>. Mean and standard deviation calculated using 5-fold cross validation. Results are reported as mean *±* standard deviation over 5-fold cross-validation.

| Model/Ablation | Features | Kuramoto (MSE) | PVC (Acc.) | fMRI (Acc.) |
| --- | --- | --- | --- | --- |
| Neurospectrum | All | **0.0041 ± 0.0002** | <u>0.60 ± 0.02</u> | <u>0.70 ± 0.02</u> |
| | BC | 0.0064 ± 0.0003 | **0.62 ± 0.01** | 0.61 ± 0.03 |
| | $\kappa$ | 0.0156 ± 0.0004 | 0.54 ± 0.02 | 0.63 ± 0.01 |
| | PS | <u>0.0044 ± 0.0002</u> | 0.59 ± 0.01 | **0.72 ± 0.01** |
| | RNN | 0.0047 ± 0.0001 | 0.53 ± 0.02 | 0.63 ± 0.02 |
| Neurospectrum-PCA | All | 0.0102 ± 0.0002 | 0.46 ± 0.03 | 0.56 ± 0.02 |
| | BC | 0.0193 ± 0.0004 | 0.39 ± 0.02 | 0.55 ± 0.03 |
| | $\kappa$ | — | — | — |
| | PS | 0.0117 ± 0.0003 | 0.44 ± 0.02 | 0.54 ± 0.02 |
| | RNN | 0.0109 ± 0.0002 | 0.43 ± 0.02 | 0.48 ± 0.03 |
| Neurospectrum-tSNE | All | 0.0143 ± 0.0012 | 0.51 ± 0.03 | 0.44 ± 0.03 |
| | BC | 0.0217 ± 0.0011 | 0.47 ± 0.02 | 0.39 ± 0.02 |
| | $\kappa$ | — | — | — |
| | PS | 0.0135 ± 0.0005 | 0.51 ± 0.02 | 0.41 ± 0.02 |
| | RNN | 0.0148 ± 0.0009 | 0.42 ± 0.02 | 0.44 ± 0.03 |
| T-PHATE | All | 0.0065 ± 0.0002 | 0.47 ± 0.02 | 0.62 ± 0.02 |
| | BC | 0.0073 ± 0.0004 | 0.46 ± 0.02 | 0.56 ± 0.02 |
| | $\kappa$ | 0.0129 ± 0.0003 | 0.36 ± 0.02 | 0.58 ± 0.03 |
| | PS | 0.0064 ± 0.0002 | 0.49 ± 0.02 | 0.65 ± 0.01 |
| | RNN | 0.0068 ± 0.0003 | 0.46 ± 0.01 | 0.61 ± 0.02 |
| GraphPH-CROCKER | | 0.0281 ± 0.0003 | 0.53 ± 0.01 | 0.52 ± 0.02 |

**Figure S1:**
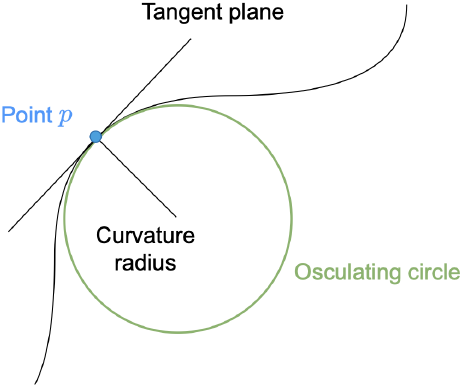
Circle fitting method for computation of curvature from latent trajectory.

**Figure S2:**
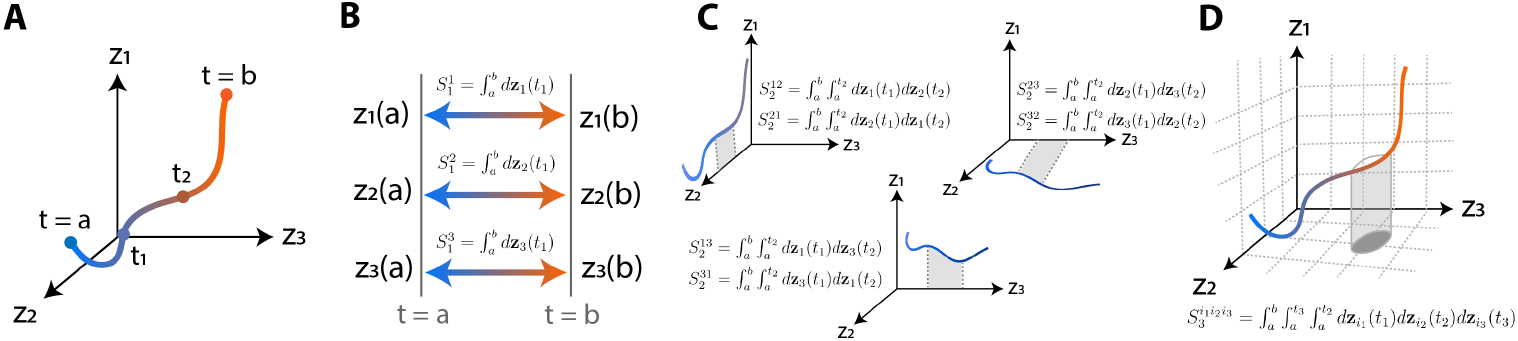
Computation of path signatures. (A) A latent trajectory **z**(*t*) in ℝ^3^, parameterized by time. Color represents temporal evolution. (B) First-order signatures *S*_1_ capture net displacement along each axis. Each component 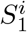 is the integral 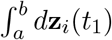. (C) Second-order signatures 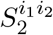 compute the signed area “swept” by the trajectory in coordinate pairs. Each term 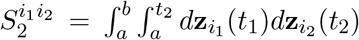 encodes curvature and directional co-dependence. (D) Third-order signatures 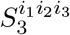 accumulate volumetric contributions from coordinated changes across three dimensions. The iterated integral 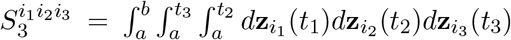 captures torsion-like effects and higher-order interactions.

**Figure S3:**
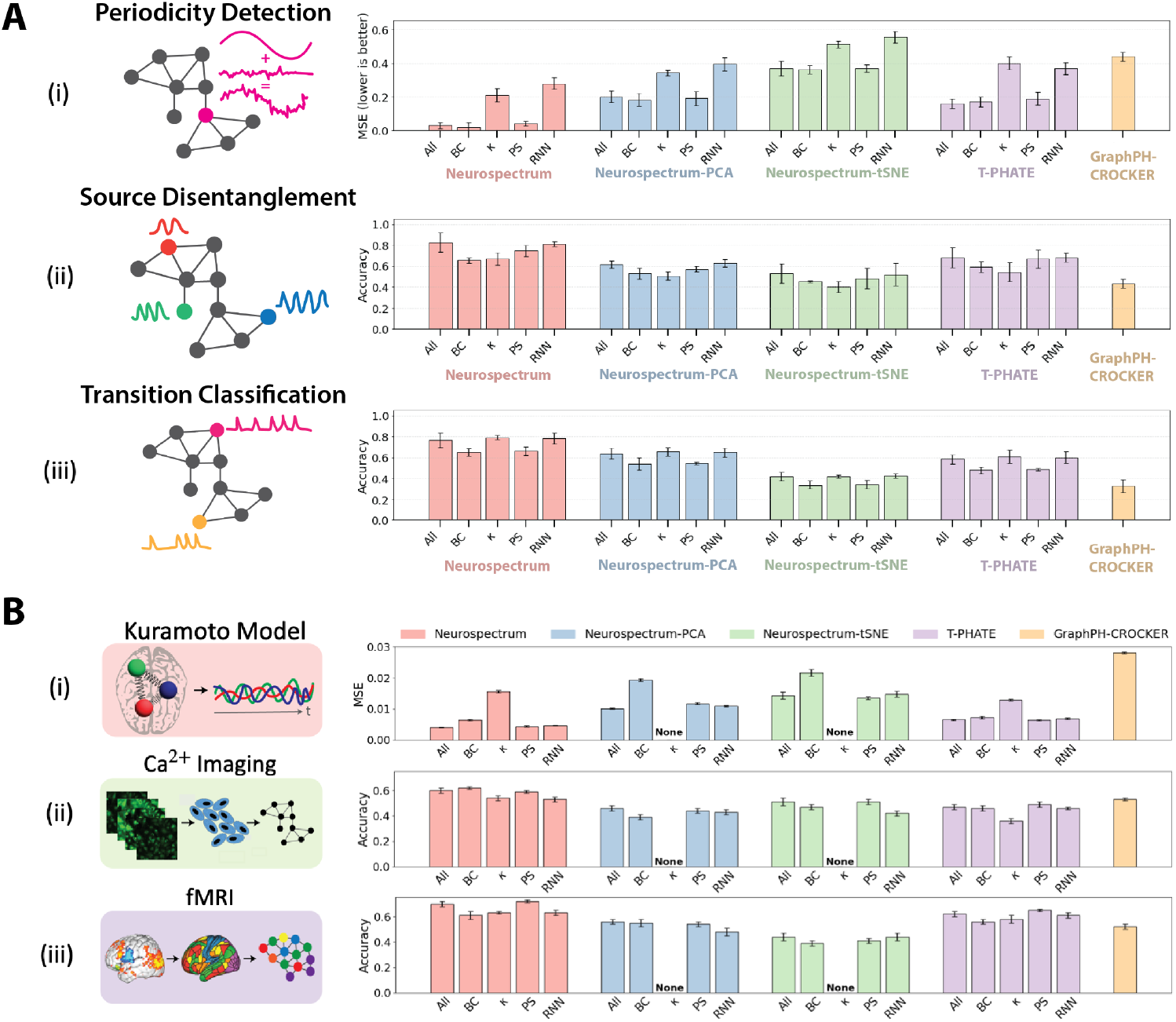
Ablation study evaluating the contributions of different latent trajectory representations in Neurospectrum. (A) Synthetic benchmarks used to assess the impact of various ablations. (i) Periodicity Detection: A sinusoidal signal is introduced at a single source node and diffused across the graph; the task is to regress its frequency. Performance is reported as mean squared error (MSE); lower is better. (ii) Source Disentanglement: Multiple sinusoidal signals with distinct frequencies originate from different graph nodes; the task is to classify the number of unique sources. Accuracy is reported; higher is better. (iii) Transition Classification: Simulated transitions from asynchronous to synchronous (collective spiking) dynamics; the task is to detect whether a transition occurred. Accuracy is reported; higher is better. (B) Real-world datasets used to assess the impact of various ablations. (i) Kuramoto Model: The task is to predict the coupling strength parameter from weakly coupled oscillator dynamics. MSE is reported; lower is better. (ii) Ca^2+^ Imaging: The task is to classify the spatial frequency and orientation of sinusoidal grating stimuli from Calcium activity in layer 4 of the mouse visual cortex. Accuracy is reported; higher is better. (iii) fMRI: The task is to distinguish OCD subjects from healthy controls using resting-state brain activity measured using fMRI. Accuracy is reported; higher is better.

**Figure S4:**
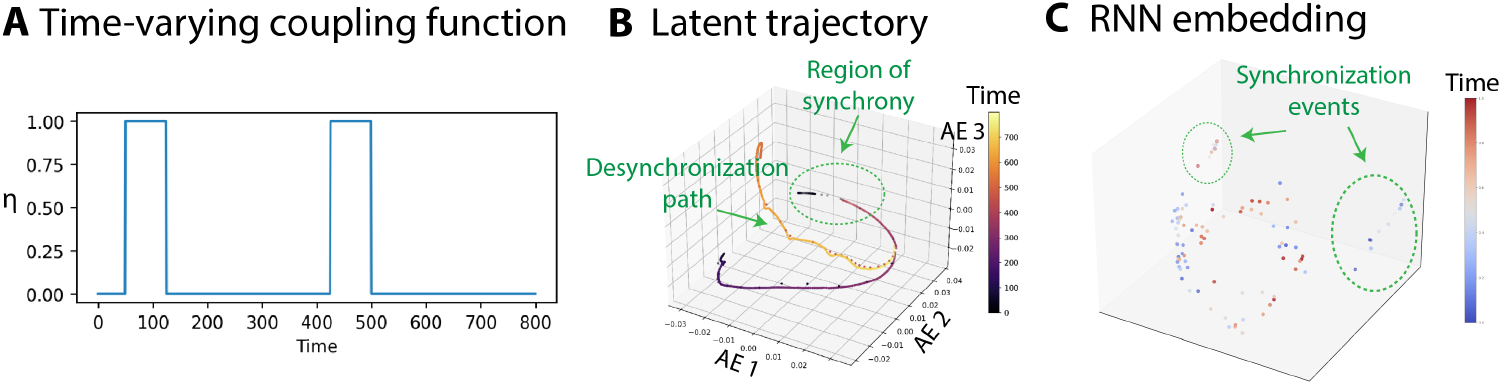
Detecting transitions in synchrony in Kuramoto model simulations of 100 coupled oscillators with a time-varying coupling strength *η*. (A) The coupling parameter *η* is initially set to 0 and increased to 1 at two distinct time windows, inducing temporary synchrony before reverting to 0. (B) The latent trajectory obtained from NeuroSPECTRUM reveals distinct dynamical regimes: timepoints corresponding to high synchrony (i.e., *η* = 1) cluster in a compact region of the latent space, while desynchronization triggered by suddenly decreasing *η* to 0 generates a smooth trajectory that re-enters this region upon resynchronization. (C) Embeddings from the final hidden state of an RNN trained to regress the value of *η* from segments of the latent trajectory show clear separation of synchronized periods. Clustering of *η* = 1 timepoints indicates that the RNN successfully learns to detect synchronization transitions from temporal patterns in the latent space.

**Figure S5:**
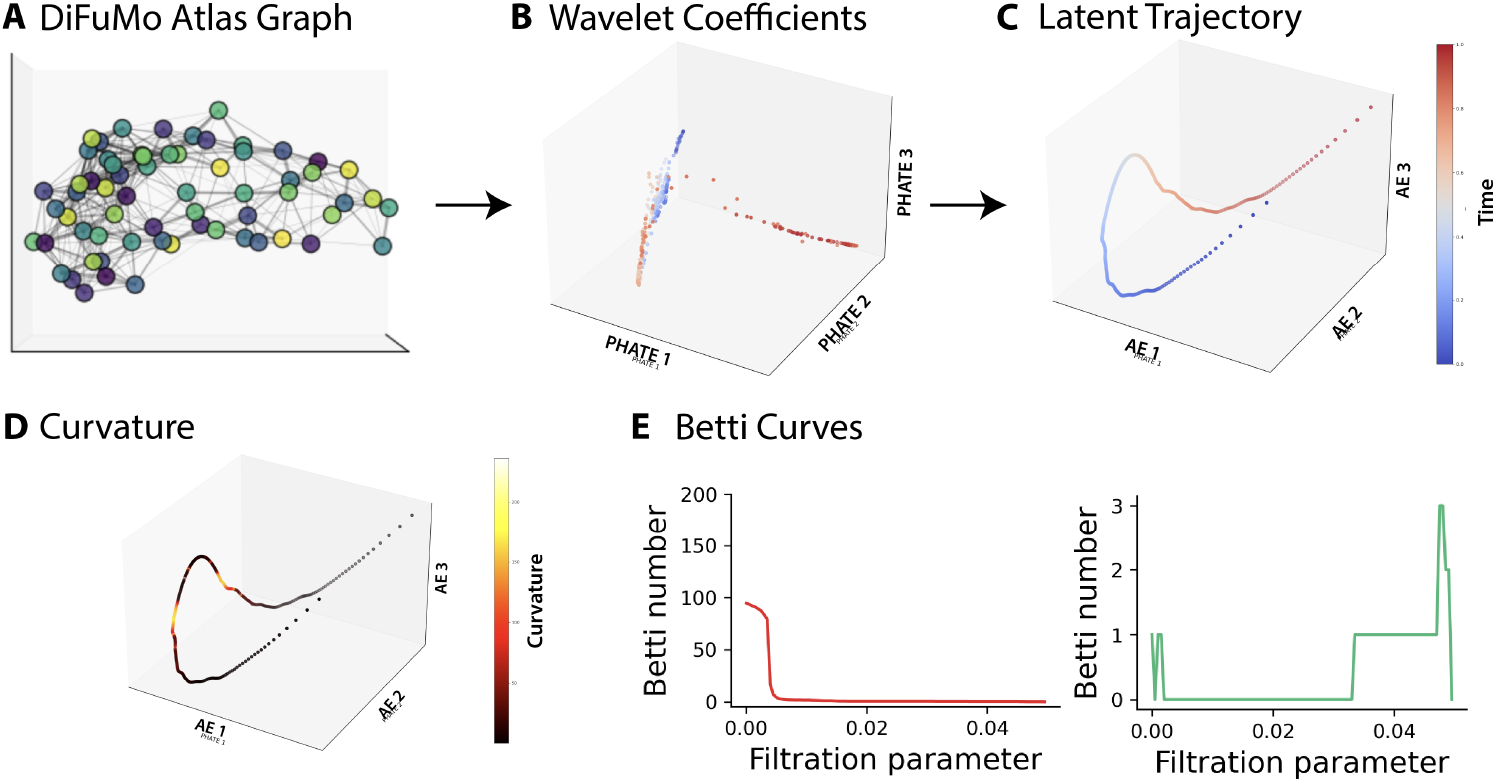
Geometrical and topological summaries of spatiotemporal dynamics of brain activity. (A) Anatomical graph constructed from the DiFuMo atlas coordinates which represents each parcel in the brain. (B) This anatomical graph is then used to obtain the wavelet coefficients via deep wavelet transform of the timelapes brain signal. Here wavelet coefficients are visualized in 3-dimensional space using PHATE. (C) The wavelet transform are subsequently used to obtain a smooth latent trajectory via the T-PHATE regularized autoencoder. (D) The curvature of the latent trajectory can be computed and is visualized in 3-dimensional space. (E) Betti curves summarize the topological features of the latent trajectory by tracking the number of connected components (left) and loops (right) across a range of filtration scales.

